# The Future of OA: A large-scale analysis projecting Open Access publication and readership

**DOI:** 10.1101/795310

**Authors:** Heather Piwowar, Jason Priem, Richard Orr

## Abstract

Understanding the growth of open access (OA) is important for deciding funder policy, subscription allocation, and infrastructure planning.

This study analyses the number of papers available as OA over time. The models includes both OA embargo data and the relative growth rates of different OA types over time, based on the OA status of 70 million journal articles published between 1950 and 2019.

The study also looks at article usage data, analyzing the proportion of views to OA articles vs views to articles which are closed access. Signal processing techniques are used to model how these viewership patterns change over time. Viewership data is based on 2.8 million uses of the Unpaywall browser extension in July 2019.

We found that Green, Gold, and Hybrid papers receive more views than their Closed or Bronze counterparts, particularly Green papers made available within a year of publication. We also found that the proportion of Green, Gold, and Hybrid articles is growing most quickly.

In 2019:

- 31% of all journal articles are available as OA
- 52% of article views are to OA articles

Given existing trends, we estimate that by 2025:

- 44% of all journal articles will be available as OA
- 70% of article views will be to OA articles

The declining relevance of closed access articles is likely to change the landscape of scholarly communication in the years to come.

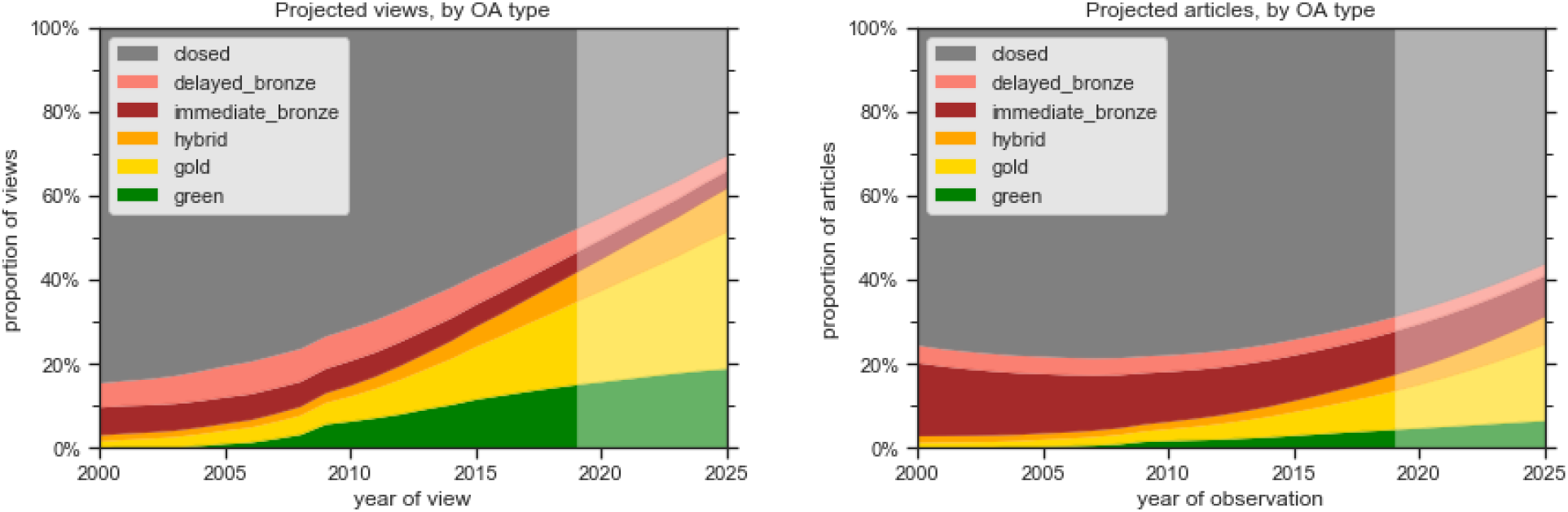

Percent of views, by OA type:

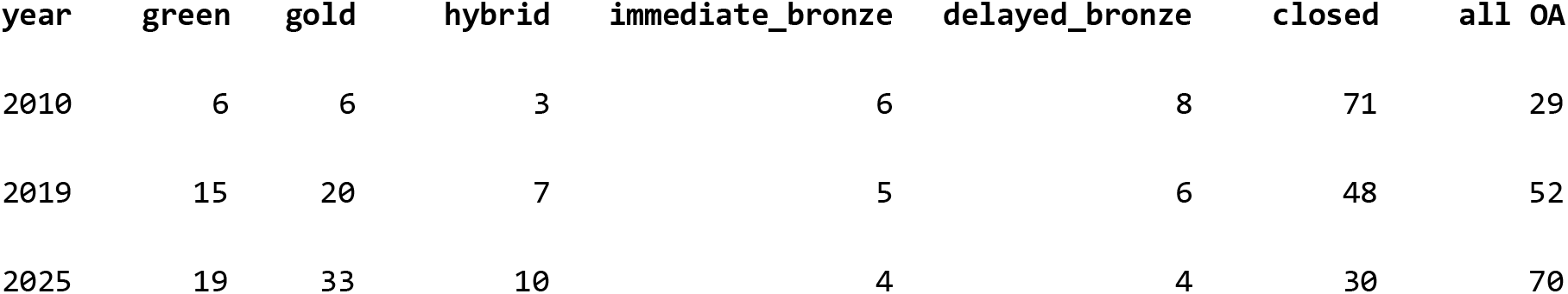

Percent of papers, by OA type:

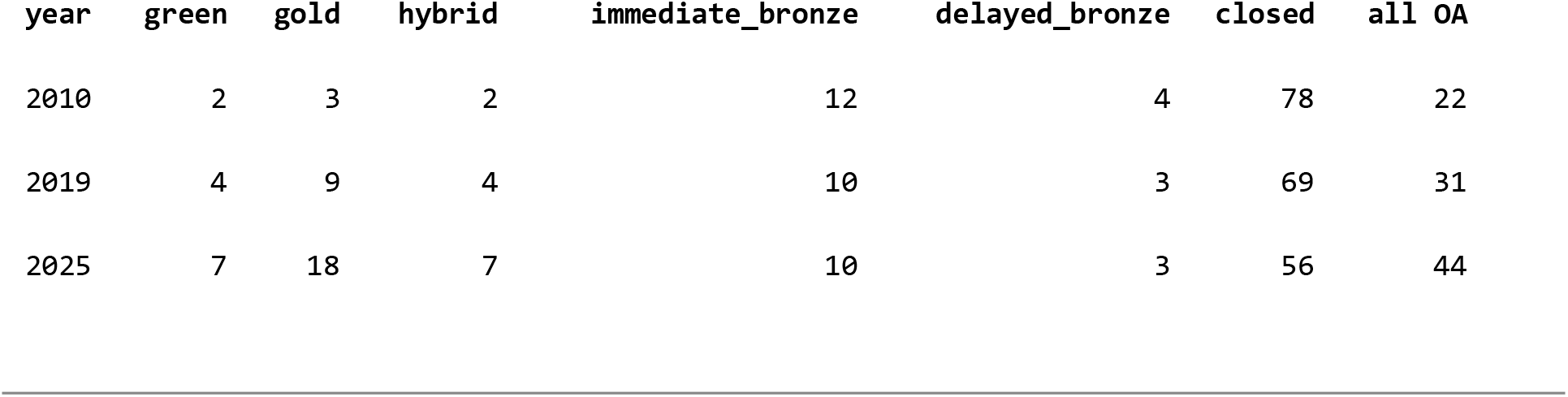

## 1. Introduction

The adoption of <u>open access (OA)</u> publishing is changing scholarly communication. Predicting the future prevalence of OA is crucial for many stakeholders making decisions now, including:

- libraries deciding which journals to subscribe to and how much they should pay
- institutions and funders deciding what mandates they should adopt, and the implications of existing mandates
- scholarly publishers deciding when to flip their business models to OA
- scholarly societies deciding how best to serve their members.

Despite how useful OA prediction would be, only a few studies have made an attempt to empirically predict open access rates. Lewis (2012) extrapolated the rate at which <u>gold OA</u> would replace subscription-based publishing using a simple log linear extrapolation of gold vs subscription market share. Antelman (2017) used one empirically-derived growth rate for <u>green</u> <u>OA</u> and another for all other kinds of OA combined. Both of these studies are based on data collected before 2012, and rely on relatively simple models. Moreover, these studies predict the number of papers that are OA. While this number is important, it is arguably less meaningful than the number of views that are OA, since this latter number describes the prevalence of OA as experienced by actual readers.

This paper aims to address this gap in the literature. In it, we build a detailed model using data extrapolated from large and up-to-date Unpaywall dataset (https://unpaywall.org/). We use the model to predict the number of articles that will be OA (including gold, green, hybrid, and bronze OA) over the next five years, and also use data from the Unpaywall browser add-on (https://unpaywall.org/products/extension) to predict the proportion of scholarly article views that will lead readers to OA articles over time.

This paper aims to provide models of OA growth, taking the following complexities into account:

- some forms of OA include a delay between when a paper is first published and when it is first freely available
- different forms of open access are being adopted at different rates
- wide-sweeping policy changes, technical improvements, or cultural changes may cause disruptions in the growth rates of OA in the future

## 2. Data

The data in this analysis comes from two sources: (1) the Unpaywall dataset and (2) the access logs of the Unpaywall web browser extension.

### 2.1 OA type: the Unpaywall dataset of OA availability

Predicting levels of open access publication in the future requires detailed, accurate, timely data. This study uses the <u>Unpaywall</u> dataset to provide this data. Unpaywall is an open source application that links every research article that has been assigned a Crossref DOI (more than 100 million in total) to the OA URLs where the paper can be read for free. It is built and maintained by Our Research (formerly Impactstory), a US-based nonprofit organization.

Unpaywall gathers data gathered from over 50,000 journals and open-access repositories from all over the world. The full Unpaywall dataset is freely, publicly available (see details: https://unpaywall.org/user-guides/research).

Our definitions of OA type (gold, green, hybrid, bronze, closed) are described in Piwowar et al. (2018). To facilitate prediction, for the purpose of this analysis we subdivided bronze OA into immediate and delayed OA. In summary, these definitions are:

- 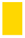 **Gold:** published in a fully-OA journal
- 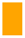 **Hybrid:** published in a toll-access journal, available on the publisher site, with an OA license
- 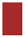 **Bronze:** published in a toll-access journal, available on the publisher site, without an OA license

- 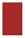 **Immediate Bronze:** available as Bronze OA immediately upon publication
- 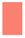 **Delayed Bronze:** available as Bronze OA after an embargo period
- 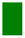 **Green:** published in a toll-access journal and the only fulltext copy available is in an OA repository
- 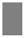 **Closed:** everything else

This analysis uses all articles with a Crossref article type of “journal-article” published between 1950 and the date of the analysis (October 2019), which is 71 million articles.

### 2.2 Article views: access logs of the Unpaywall web browser extension

Predicting the open access pattern of usage requests requires knowing the relative usage demands of papers based on their age. This study has extracted these pageview patterns from the usage logs of the <u>Unpaywall browser extension</u> for Chrome and Firefox.

This extension is an open-source tool made by the same non-profit as the Unpaywall dataset described above, with the goal of helping people conveniently find free copies of research papers directly from their web browser. The extension has <u>more than 200,000 active users</u>, distributed globally, as shown in Figure 1.

**Figure 1:**
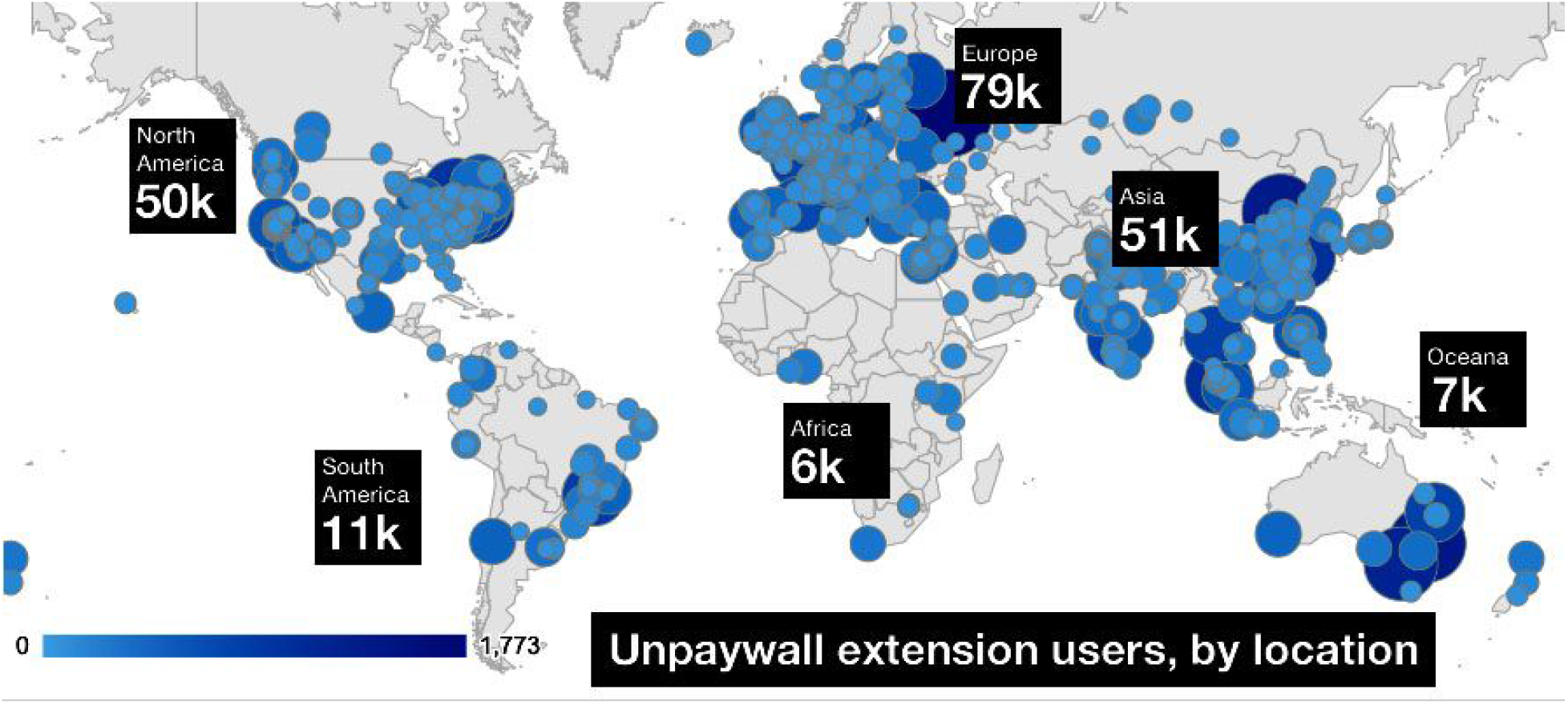
Map of Unpaywall users in February 2019.

The Unpaywall browser automatically extension detects when a user is on a scholarly article webpage -- we consider this an access request, or a view. The extension can be disabled, or can be configured to only run upon request, but very few users use these settings.

The extension received more than 3 million article access requests in July 2019 which we use for most of our analysis. Because readership data is private and potentially sensitive, we are not releasing the Unpaywall usage logs along with the other datasets behind this paper other than as aggregate counts by OA type by year.

## 3. Approach

### 3.1 Overview

The goal of this analysis is to predict two aspects of OA growth:

1. Growth in OA articles and their proportion of the literature over time
2. Growth in OA article views and their proportion of all literature views over time

We examine the growth in OA articles *by date of observation*, rather than by date of publication. This requires us to calculate the OA lag between publication and availability for different types of OA, which is done in Section 4.1.

Once we have the pattern of OA availability by year, we forecast the OA availability for future years by assuming that it will have the same overall pattern as previous years -- the papers that will be made available next year will have the same age distribution as papers that were made available last year. We allow the absolute number of papers to increase year-over-year: we estimate the future growth multiplier by extrapolating the height of past availability curves. This analysis is presented in Section 4.2.

Next, we turn to predicting the growth of OA article views -- what proportion of what is read is available OA, and how will this change in the future? The Unpaywall browser extension logs give us a relative baseline of what is read right now. By assuming that reading patterns remain relatively unchanged over time (specifically the probability that a reader wants to read a paper given its age and OA type), we use the publication estimates we made in previous sections to calculate the relative number of views by OA type in the past and the future. This is described in Section 4.3.

Finally, we look at the impact of extending the model to include a disruptive change, in this case the growth of bioRxiv, in Section 4.4.

### 3.2 Glossary

In addition to the OA types defined in Section 2.1, we define additional terms as we use them in this paper, in approximate order they are discussed:

- **Date of publication**: the date an article is published in a journal
- **Embargo**: the delay that some toll-access journals require between date of publication and when an article can be made Green or Delayed Bronze OA
- **Self-archiving**: when an author posts their article in an OA repository
- **OA type**: the OA classification of an article, as defined in Section 2. The OA type of an article may change over time (from Closed to Delayed Bronze OA, or from Closed to Green OA) because of embargoes and other self-archiving delays
- **Date first available OA**: the date an article first becomes an OA type other than “Closed”
- **OA lag**: the length of time between an article’s Date of Publication and its Date First Available OA
- **OA assessment**: the determination of the OA type of an article at a given point in time
- **Date of observation**: the point in time for which we make an OA assessment for an article. Explained in Section 3.3.
- **Observation age** of an article: the length of time between an OA assessment observation and the article’s date of publication
- **View**: someone on the internet visited the publisher webpage of an article, presumably with the hope of reading the article
- **Date of view**: the date of the view
- **View age** of an article: the length of time between an article’s date of publication and the date of a view
- **Articles by age curve**: for a given snapshot, the plot of snapshot age (in years) on the x-axis and number of articles published of that snapshot age on the y-axis
- **Views by age curve**: the plot of view age (in years) on the x-axis and number of views received by articles of that view age on the y-axis
- **Views per article by age curve**: the plot of view or snapshot age (in years) on the x-axis and number of views per article (by views of that view age and articles of that snapshot age) on the y-axis
- **Views per year curve**: the plot of year on the x-axis and the number of views estimated to have been made that year on the y-axis

### 3.3 Date of Observation

In this paper we approach the growth of OA from the Date of Observation of OA assessment, rather than the date of publication. We explain this with the use of Figure 2.

**Figure 2:**
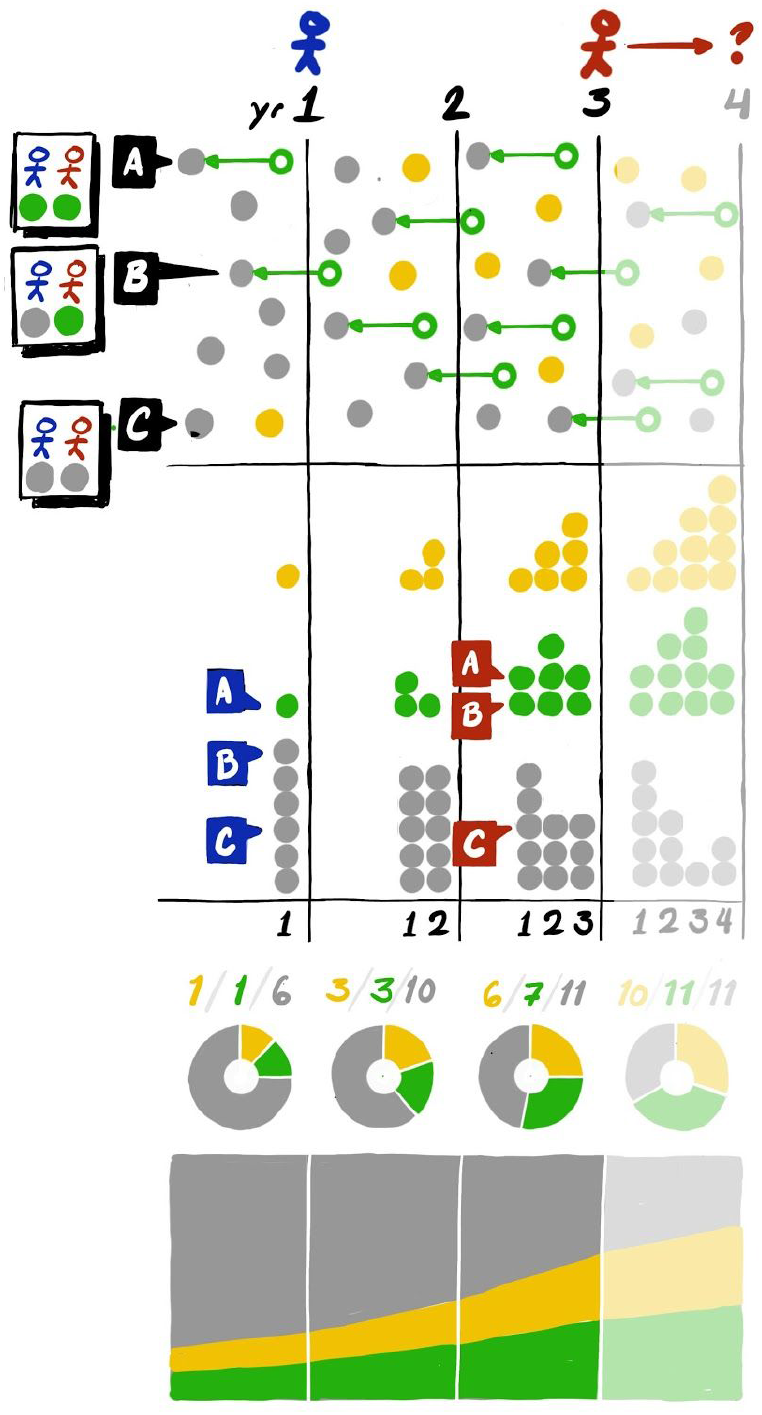
Date of observation.

Let’s imagine two observers, Alice (blue) and Bob (red), shown by the two stick figures at the top of Figure 2.

Alice lives at the end of Year 1--that’s her “Date Of Observation.” Looking down, she can see all 8 articles (represented by solid colored dots) published in Year 1, along with their access status: Gold OA, Green OA, or Closed. The Year of Publication for all eight of these articles is Year 1.

Alice likes reading articles, so she decides to read all eight Year 1 articles, one by one.

She starts with Article A. This article started its life early in the year as Closed. Later that year, though--after an OA Lag of about six months--Article A became Green OA as its author deposited a manuscript (the green circle) in their institutional repository. Now, at Alice’s Date of Observation, it’s open! Excellent. Since Alice is inclined toward organization, she puts Article A article in a stack of Green articles she’s keeping below.

Now let’s look at Bob. Bob lives in Alice’s future, in Year 3 (ie, his “Date of Observation” is Year 3). Like Alice, he’s happy to discover that Article A is open. He puts it in his stack of Green OA articles, which he’s further organized by date of their publication (it goes in the Year 1 stack).

Next, Alice and Bob come to Article B, which is a tricky one. Alice is sad: she can’t read the article, and places it in her Closed stack. Unbeknownst to poor Alice, she is a victim of OA Lag, since Article B will become OA in Year 2. By contrast, Bob, from his comfortable perch in the future, is able to read the article. He places it in his Green Year 1 stack. He now has two articles in this stack, since he’s found two Green OA articles in Year 1.

Finally, Alice and Bob both find Article C is closed, and place it in the closed stack for Year 1. We can model this behavior for a hypothetical reader at each year of observation, giving us their view on the world--and that’s exactly the approach we take in this paper.

Now, let’s say that Bob has decided he’s going to figure out what OA will look like in Year 4. He starts with Gold. This is easy, since Gold article are open immediately upon publication, and publication date is easy to find from article metadata. So, he figures out how many articles were Gold for Alice (1), how many in Year 2 (3), and how many in his own Year 3 (6). Then he computes percentages, and graphs them out using the stacked area chart at the bottom of Figure 2. From there, it’s easy to extrapolate forward a year.

For Green, he does the same thing--but he makes sure to account for OA Lag. Bob is trying to draw a picture of the world every year, as it appeared to the denizens of that world. He wants Alice’s world as it appeared to Alice, and the same for Year 2, and so on. So he includes OA Lag in his calculations for Green OA, in addition to publication year. Once he has a good picture from each Date Of Observation, and a good understanding of what the OA Lag looks like, he can once again extrapolate to find Year 4 numbers.

Bob is using the same approach we will use in this paper--although in practice, we will find it to be rather more complex, due to varying lengths of OA Lag, additional colors, of OA, and a lack of stick figures.

### 3.4 Statistical analysis

The analysis was implemented as an executable python Jupyter notebook using the pandas, scipy, matplotlib, and sqlalchemy libraries. See the Data and code availability section below for links to the analysis code and raw data.

## 4. Methods and Results

### 4.1 Past OA Publication, by date of observation

#### 4.1.1 OA lag

For Gold OA and Hybrid OA understanding OA lag is easy -- there is no lag: papers become OA at the time of publication.

For Green and Bronze OA the lag is more complicated. Authors often self-archive (upload their paper to a repository) months or years after the official publication date of the paper, typically because the journal has a policy that authors must wait a certain length of time (the “embargo period”) before self-archiving. Funder policies that mandate Green OA often allow a delay between publication and availability (notably the National Institutes of Health in the USA allows a 12 month embargo, which is relevant for most of the content in the large PMC repository). Finally, some journals open up their back catalogs once articles reach a certain age, which has been called “delayed OA” (Laakso and Björk, 2013) and we consider an important subset of Bronze.

We explore and model these dynamics below.

#### 4.1.2. OA lag for Green OA

Calculating OA lag requires data on both when an article was first published in its journal and the date it was first made OA.

The date an article becomes Green OA can be derived from the date it was made available in a repository, which we can get from its matched <u>OAI-PMH records</u> (as harvested by Unpaywall).

Figure 3 shows four plots: the leftmost plot shows Green OA articles that were first made OA in 2015, the second plot shows Green OA articles that were first made OA in 2016, and so on. Each plot is a histogram of number of articles by date of publication.

**Figure 3:**
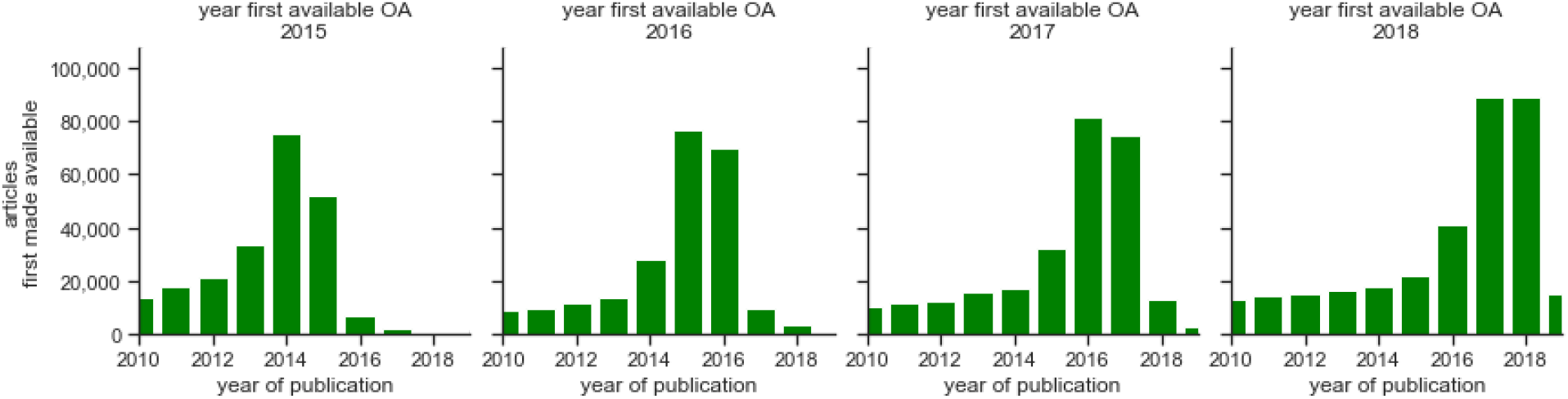
OA lag for Green OA. Each plot shows articles that were first made available during the given year of observation, by year of their publication on the x-axis.

By looking at the first plot in depth, we can see that a few articles are made available *before* they are actually published (articles published in 2016 or 2017) -- these were preprints, submitted before publication. Continuing to look at the first row, we can see the bulk of the articles that became available in 2015 were published in 2015 (lag of zero years) or in 2014 (lag of 1 year). A few were published in 2013 (an OA lag of 2 years), and then a long tail represents the backfilling of older articles.

Looking now at all plots in Figure 3, we can see that a similar OA lag pattern (a few preprints are available before publication, most articles become available within a 3 year OA lag, then a long tail) has held for the last four years of Green OA availability (the distribution of the bars are similar in all four graphs).

We can also see that the relative amount of green OA is growing slightly by year of OA-first-availability (the area under the whole histogram gets higher with subsequent histograms). Green OA appears to be growing. We will explore this further in Section 4.2.

More details on Green OA lag are included in Supplementary Information, Section 11.1.

#### 4.1.3 OA lag for Bronze Delayed OA

There was no recent, complete, publicly-available list of Delayed OA journals, so we derived a list empirically based on the Unpaywall database. We have made our list publicly available: details are in Section 7.2.

To create the list we started by looking at existing compilations of Delayed OA journals, including:

- https://www.elsevier.com/about/open-science/open-access/open-archive
- http://highwire.stanford.edu/cgi/journalinfo#loc
- https://www.ncbi.nlm.nih.gov/pmc/journals/?filter=t3&titles=current&search=journals#csvfile
- https://en.wikipedia.org/wiki/Category:Delayed_open_access_journals
- Delayed open access: An overlooked high-impact category of openly available scientific literature by Laakso and Björk (2013).

From those sources we determined that almost all embargoes for Delayed OA journals are at 6, 18, 24, 36, 48, or 60 months.

Next we used the Unpaywall data to calculate the OA rate of all journals, partitioned by age of their articles. We looked at Bronze OA rates before and after each of these common month cutoffs, highlighting cutoffs where OA was much less than 90% before the cutoff and 90% or higher afterwards. For each cutoff that looked like a Delayed OA candidate, we manually examined the full OA pattern for the journal and made a judgment call about whether it had an OA pattern consistent with a Delayed OA journal (low OA rates for articles until an embargo date, then high OA rates). We finally cross-referenced this empirically derived list with the sources again to see if it was roughly equivalent for journals on both lists -- it is, and the empirically derived list is more comprehensive.

Our resulting list includes 3.6 million articles (4.9% of all articles) published in 546 journals, with the following embargo lengths:

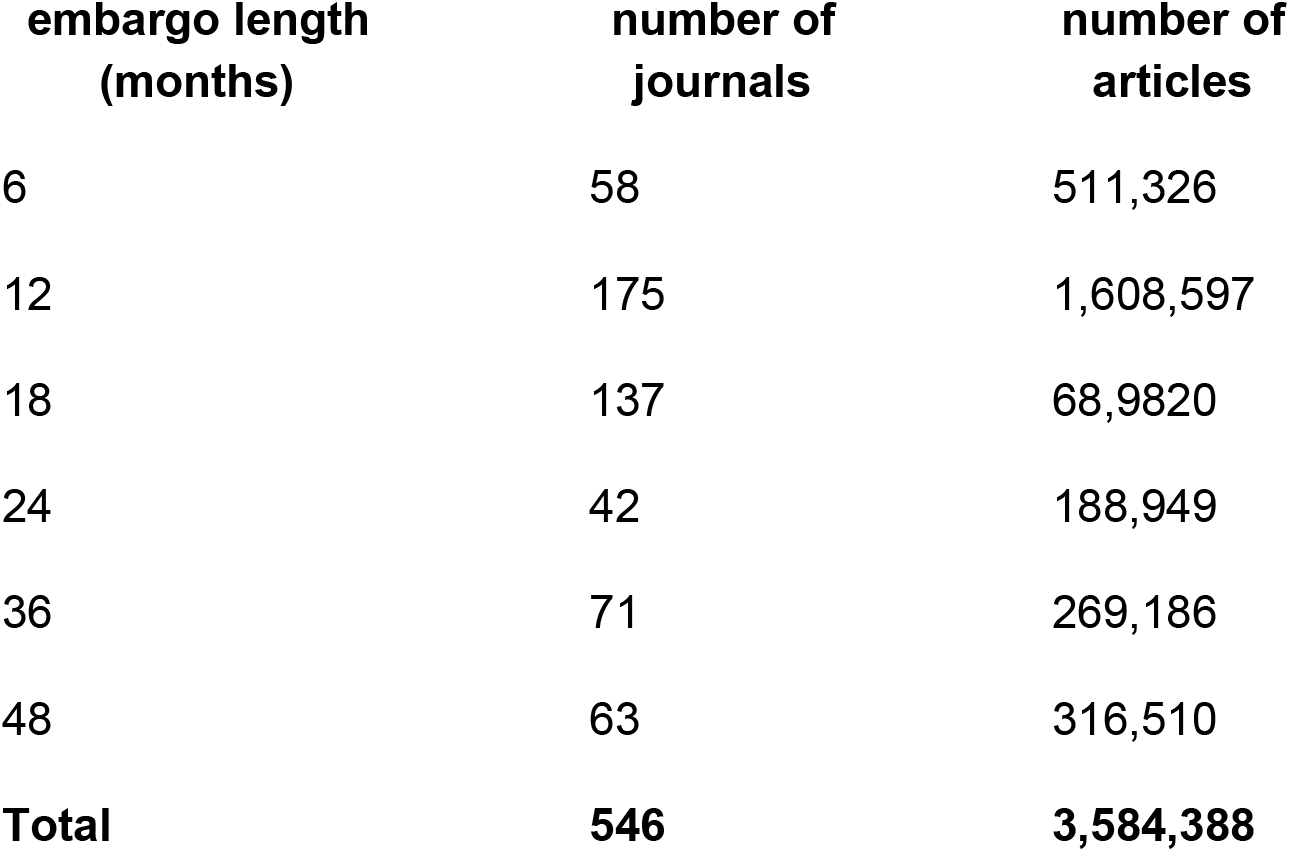

We used this list to split articles labelled “Bronze” by Unpaywall into two categories: “Delayed Bronze” for articles published in journals in our Delayed OA list, and “Immediate Bronze” for all others.

Immediate Bronze articles have no OA lag: they become available on the publisher site immediately.

We estimate the OA lag for a Delayed Bronze OA article as the Delayed OA embargo for journal it is published in. From there we can also estimate the date it first became OA by adding the embargo period to the publication date of the article.

Figure 4 shows four plots: the leftmost plot shows Delayed Bronze OA articles that were first made OA in 2015, the second plot shows Delayed Bronze OA articles that were first made OA in 2016, and so on. Each plot is a histogram of number of articles by date of publication.

**Figure 4:**
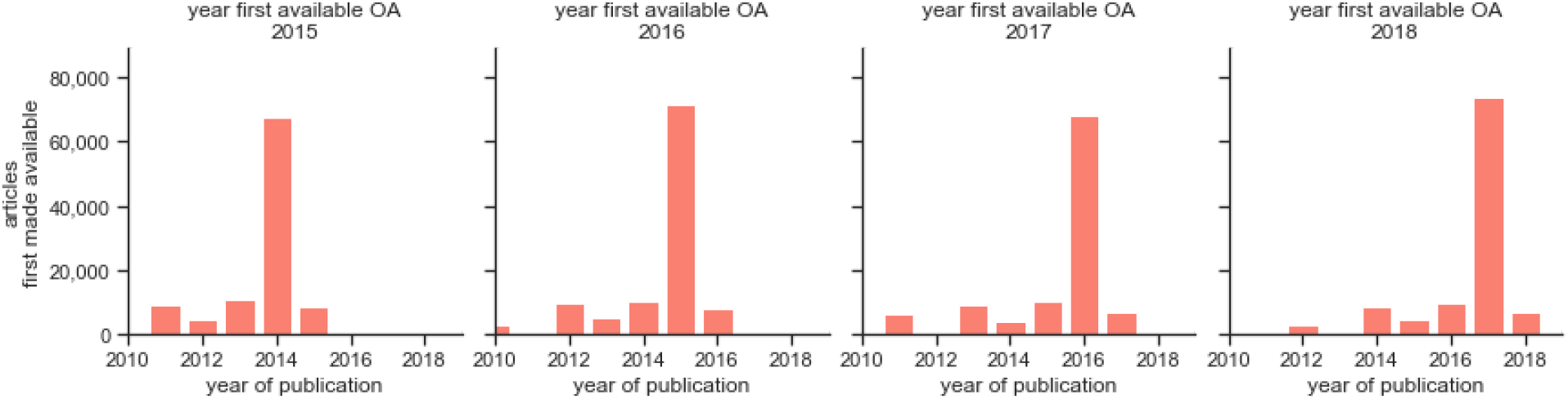
OA lag for Delayed Bronze OA. Each plot shows articles that were first made available during the given year of observation, by year of their publication on the x-axis.

The distribution of Delayed Bronze OA articles by date first made OA is shown in Figure 4, as histograms by publication date. Most articles become available after a 1 year lag. Bumps that represent articles that become available at 24, 36, and 48 months are also clearly visible.

By looking at the first plot of Figure 4 in depth, we can see that most articles first made available in Delayed Bronze OA journals were made available with 1 year OA lag, in 2014. A few were made available with a lag of less than one year, 2 years, or 4 years.

We can also see that the relative amount of Delayed Bronze OA is not growing very much by year of OA-first-availability (the area under the whole histogram gets higher with subsequent histograms is approximately the same for all histograms). Delayed Bronze OA is not growing quickly. We will explore this further in Section 4.2.

More details on Delayed Bronze OA lag are included in Supplementary Information, Section 11.2.

#### 4.1.4 Closed access at date of observation

We consider an article Closed if it has been published and is not considered OA at the time of observation.

#### 4.1.5 Past OA by date of observation and date of publication

We combine the OA lag data above to describe OA by date of observation for all OA types, in Figure 5.

**Figure 5:**
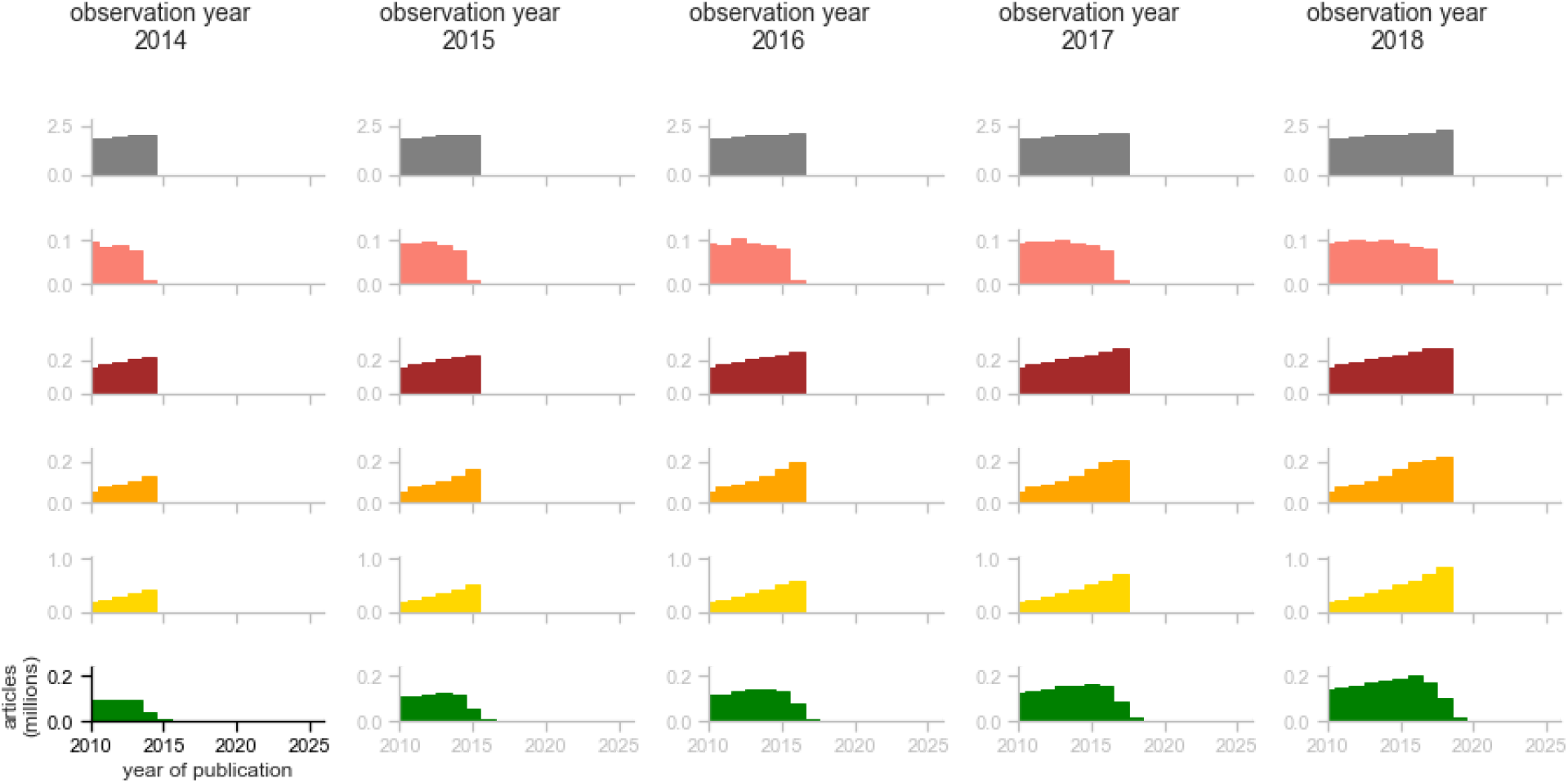
Articles by year of observation, 2014-2018. Each row is an OA Type, each column is a Year of Observation, the x-axis of each graph is the Year of Publication, and the y-axis is the total number of articles (millions) available at the year of observation.

Each column is a year of observation, from 2014 to 2018. Each row is a different OA type. Each mini plot is a histogram of all articles available by publication date, for the given observation year and OA type.

This figure differs from Figure 3 and Figure 4 in that it is cumulative over date of first availability: it shows all papers published prior to the year of observation.

We can see that Gold, Hybrid, and Immediate Bronze OA articles all simply accumulate new articles each year, immediately. For example, the 2015 Gold graph is identical to the 2014 Gold graph beside it, other than the addition of a new, taller rightmost bar showing new papers published and made available in 2015.

In contrast, Green OA (6th row) and Delayed Bronze OA (2nd row) graphs all have more complicated trends. The graphs for the 2015 observation year differ from the 2014 graphs beside them in that they have a few new publications in 2015, but they also boost the 2014 publication year, and even older years. In fact we can see that when observed in 2018 (the last column of the whole figure) Green OA is higher in all publication years than it was in the observation year 2014 (the first column in the figure) because of met embargoes and backfilling. A similar trend is visible for Delayed Bronze OA.

It is hard to see at the scale of Figure 5, but the Closed access graphs (top row) have the opposite trend -- when observed in 2018 (the last column), *fewer* papers in early bars of the histogram were considered Closed OA compared to an observation made in 2014 (first column). This is because some of what was “observed” as Closed in 2014 has become Green and Bronze by the observation year of 2018, and therefore no longer appears in the Closed access histograms.

#### 4.1.6 Combined Past OA by date of observation

We can now graph all papers by OA availability, by taking the area under each histogram in Figure 5. This gives us Figure 6, with the absolute number of articles on the left panel and proportion by OA type on the right panel. We can see that the number of OA articles has been growing over time between 2000 and 2018, though slowly. Looking at the last year of observation in the proportion graph and its table below shows 27% of the literature published by 2018 could be observed as OA in 2018.

**Figure 11:**
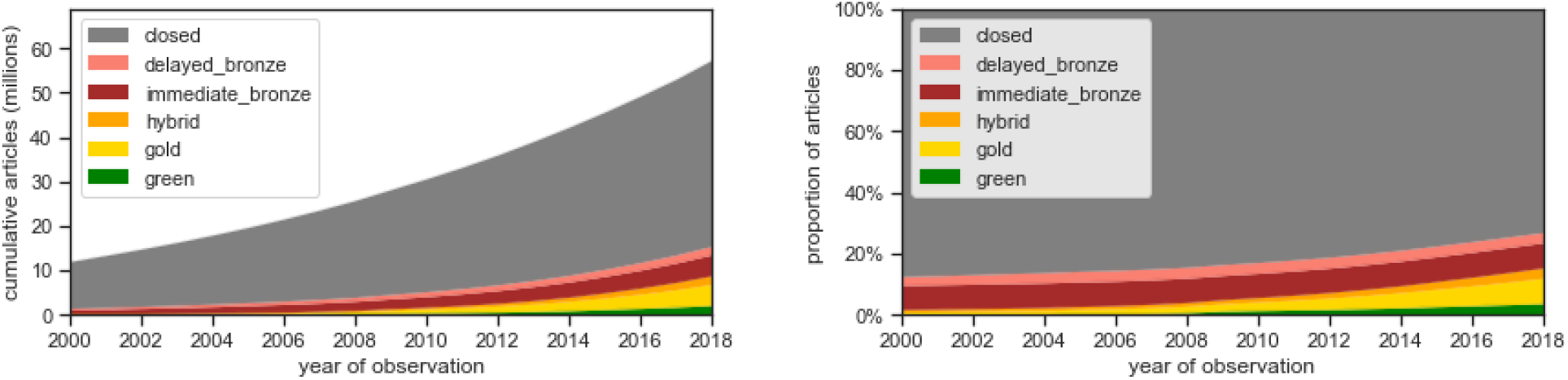
Total articles by OA type, by year of observation. OA type as of year of observation.

Table of percentages for the right panel:

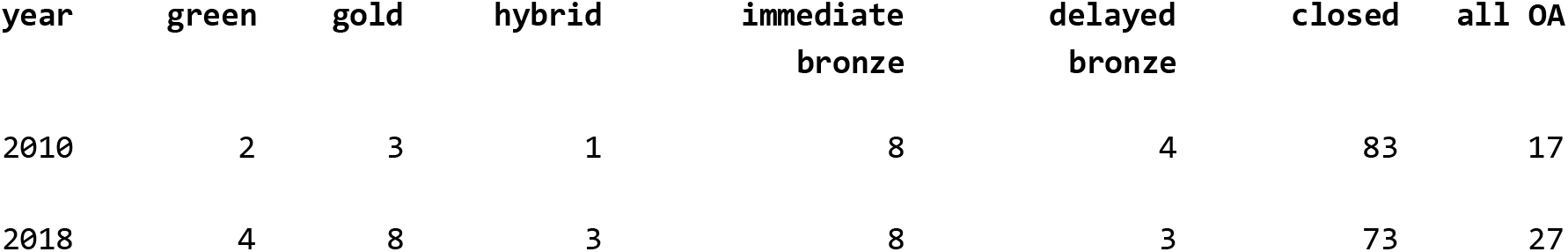

### 4.2 Future OA Publication, by date of observation

#### 4.2.1 Approach

We wish to project OA availability Figure 5 in future years. How can we extrapolate these graphs into the future?

The model we use is based on observing that the papers that become available each year have a consistent distribution by article age, as seen in Figure 3 for Green OA and Figure 4 for Bronze OA (the histograms within each figure have a similar shape).

If we then assume that the articles that will become available next year are similar to the articles that became available this year, for a given article age and OA type we can predict the future like this:

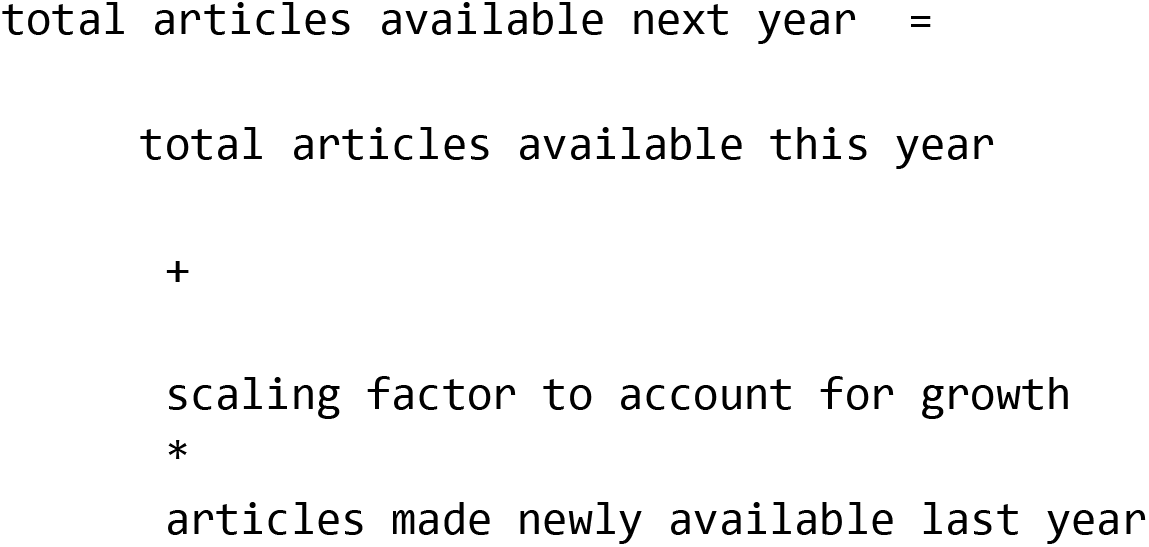

We have much of what we need already calculated in previous sections: the **total articles available this year** is the observation year 2018 in Figure 5, and the **articles made newly available last year** is the last histogram of Figure 3 for Green OA and Figure 4 for Bronze OA.

All that remains is to calculate the **scaling factor to account for growth**. We do this next.

#### 4.2.2 Scaling factor

Figure 7 shows a scatter plot of new articles by OA type, by year of observation. We add a linear best fit line, using the scipy.optimize.curve_fit() function. The r^2^ value below each graph is the sum of squares between the data and the fit, indicating goodness of fit (close to 1.0 is better).

**Figure 7:**
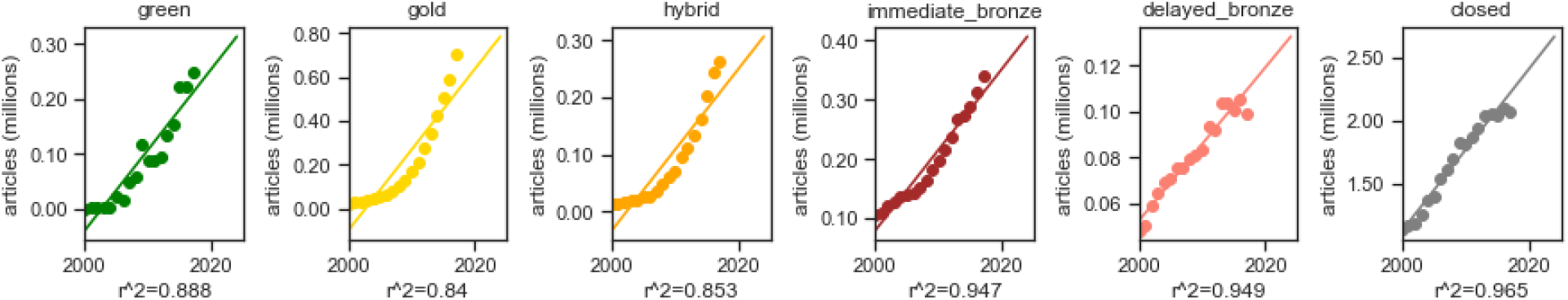
Total articles by year of observation, by OA type, with a linear extrapolation.

We can see this isn’t a particularly good fit for any of the OA types, so we try fitting with an exponential curve e^x^ in Figure 8. This is a better fit for the first four OA types, which we can see both visually and because they have higher r^2^ values.

**Figure 8:**
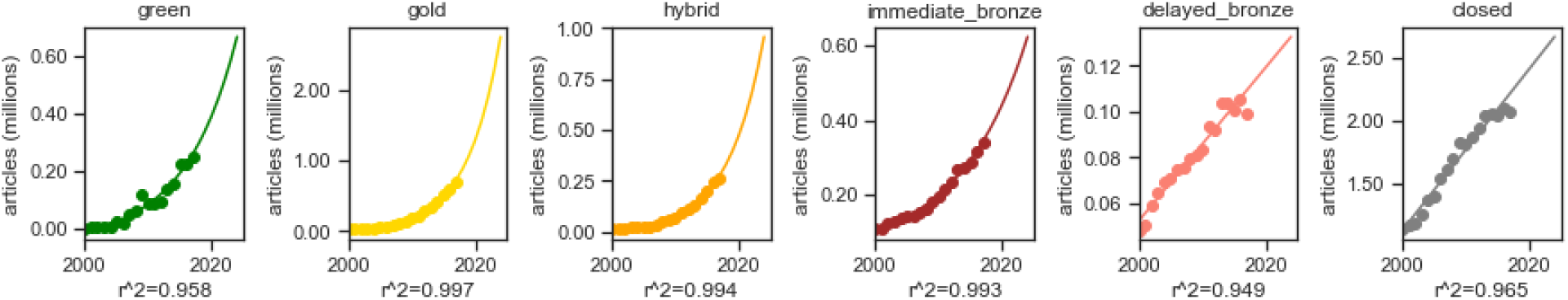
Total articles by year of observation, by OA type, with an exponential extrapolation.

The Delayed Bronze and Closed data look like they may actually trend down, so something of the form 1 - e^x^ may be a better fit. This is shown in Figure 9 for all OA Types and does indeed look like the best fit yet for both Delayed Bronze and Closed.

**Figure 9:**
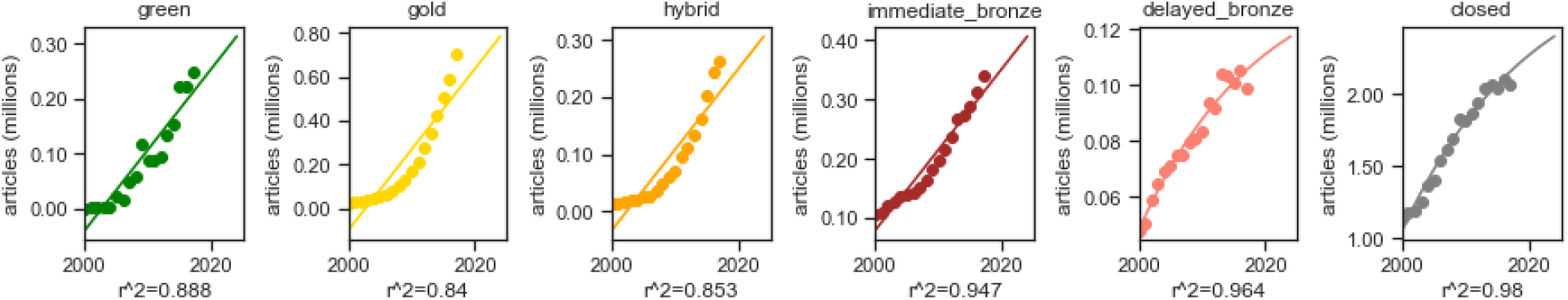
Total articles by year of observation, by OA type, with an exponential extrapolation fitting 1-exp().

We conclude this hunt for the best scaling factors by choosing the extrapolation function with the highest r^2^ value for each OA Type. We use the chosen curves to extrapolate through 2025, and use the ratio of the value in 2018 to each subsequent observation year as that year’s scaling factor.

#### 4.2.3 Future OA by date of observation, by date of publication

We now have all the information we need to calculate **total articles available next observation year** as described in Section 4.2.1. We use this approach to calculate total articles available at observation year 2019 based on 2018 data, then apply it again to calculate total articles at observation year 2020, and so on until 2025. The result is shown in Figure 10.

**Figure 10:**
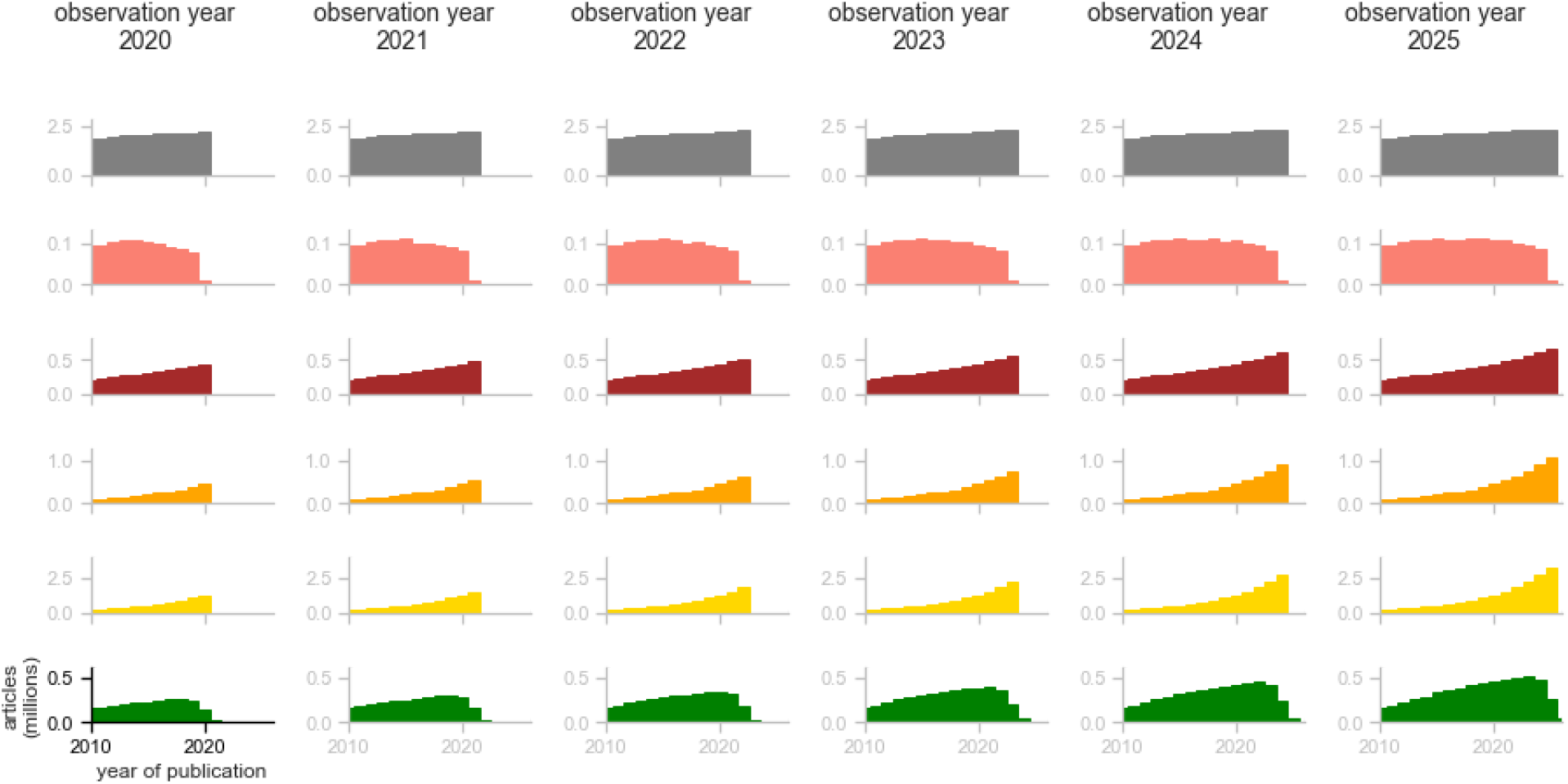
Articles by year of observation, extrapolated into the future. Each row is an OA Type, each column is a Year of Observation, the x-axis of each graph is the Year of Publication, and the y-axis is the total number of articles (millions) available at the year of observation.

#### 4.2.4 Combined Future OA by date of observation

Finally, as in Section 4.1.6, we can sum the area under the histograms above to calculate total articles by year of observation by OA Type, for past articles as well as future projections. We show this in Figure 11.

**Figure 11:**
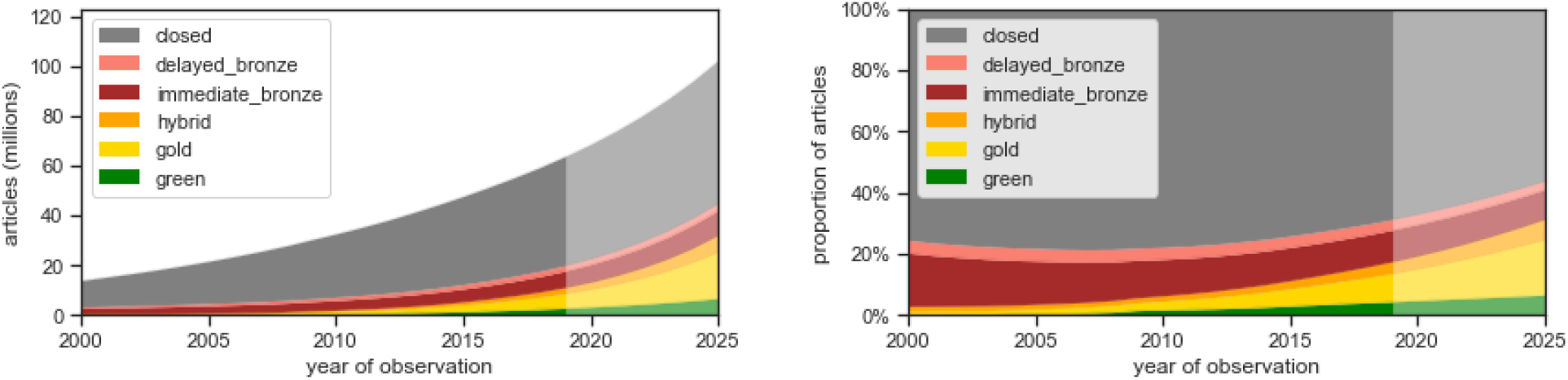
Total articles by OA type, by year of observation. OA type as of year of observation.

We project 44% of articles will be OA by 2025: Gold will account for 15% of all articles, Bronze 13%, and Green and Hybrid 7% each. A table showing the proportions of the right panel is below:

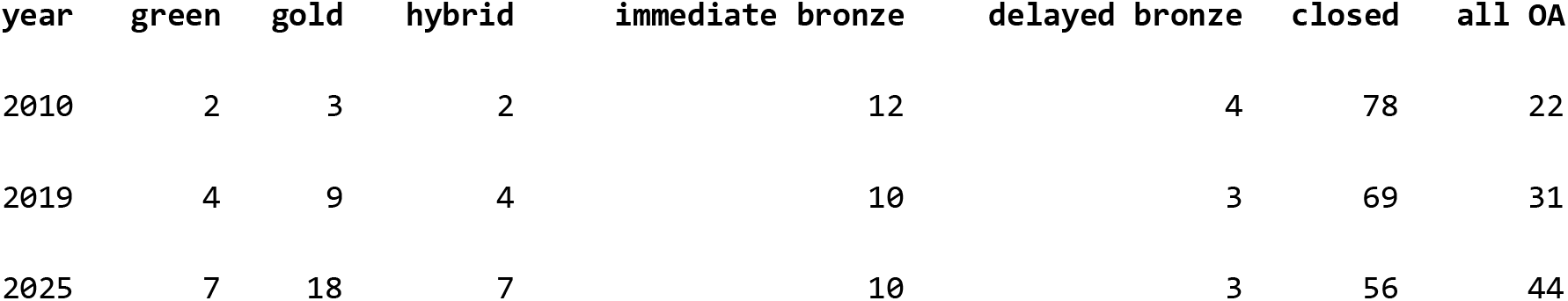

If we plot the difference between observation years in Figure 11, we get the *net change* in articles by OA type, by year of observation. This net change is shown in Figure 12.

**Figure 12:**
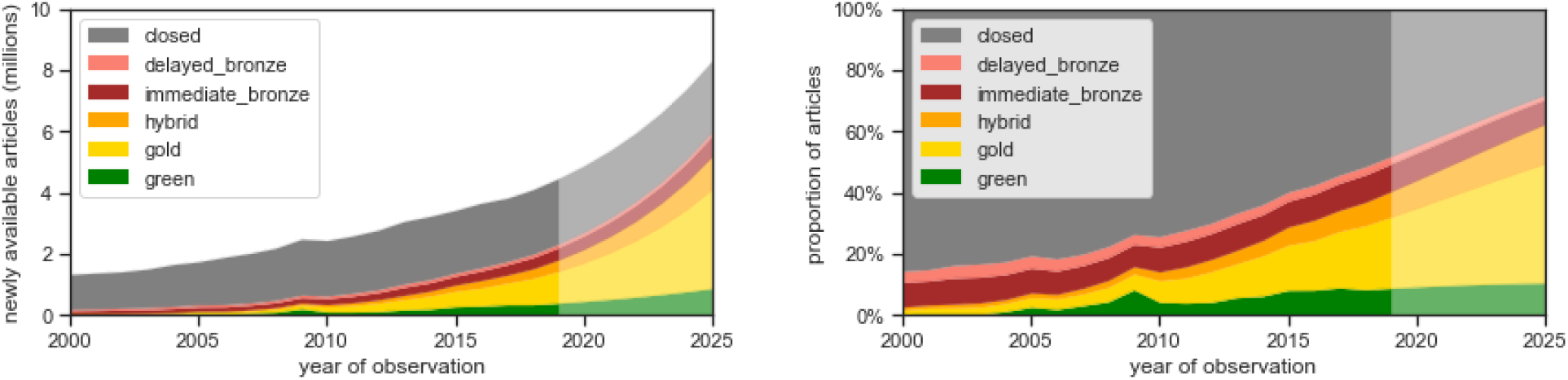
Change in number of articles from previous year of observation, by OA type. Includes newly published articles, as well as articles that have changed OA type.

This shows that by 2025 72% of articles that are newly available every year are available as OA, compared to 52% in 2019. About half of the articles that become OA each year are Gold.

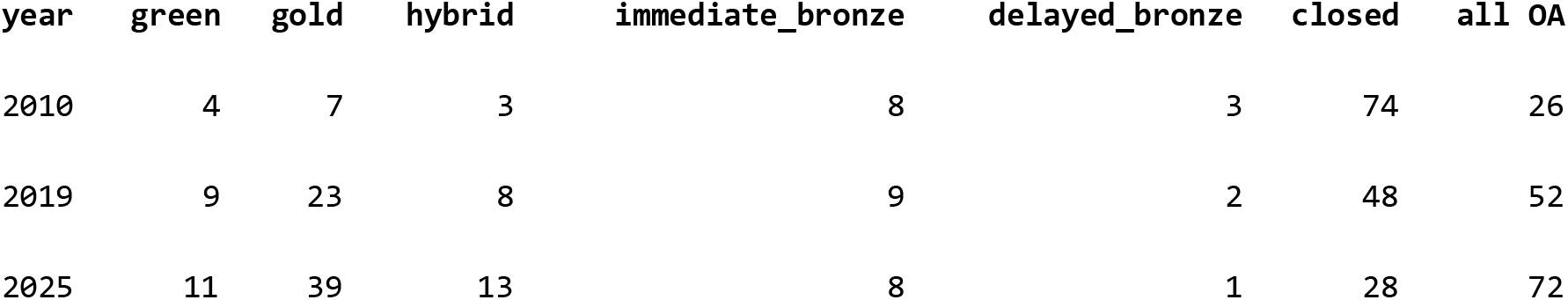

### 4.3 Past and Future OA Views

#### 4.3.1 Approach

Now that we have projections of publication trends, we change tack to examine *views* -- when people access the literature, how likely is it the article they want to read is available as OA?

How do we think this has changed over the years, and what patterns do we project in the future?

To answer these questions we will use data from the Unpaywall browser extension, as described in Section 2.2 above. This data allows us to make inferences about overall readership trends.

We will estimate views using this general equation:

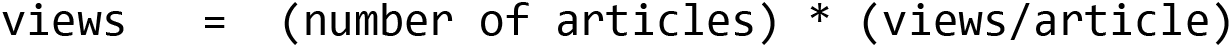

A key assumption underlying our model is that the **views/article by age of article** distribution curve is stable over time, for each OA type. We calculate this distribution for views made during July 2018, and assume that readers in all other months and years, past and future, will have a similar relative interest in articles based on their age and OA type -- we assume the number of views varies solely based on number of articles available of each age and OA type.

Because we want to know views over time (rather than just at a single point in time) we treat each of the terms in the above equation as a <u>signal</u> and use <u>digital signal processing</u> calculation techniques, which we will describe as we encounter them and in supplementary information.

The signals for **number of articles** were already calculated in Sections 4.1 and 4.2. The signals for **views/article** will be calculated in Section 4.3.2.

We “multiply” these two signals together using signal processing techniques, described in Section 4.3.3, to get total views across time.

We do these calculations for each OA type individually, and then add them together in Section 4.3.5 to look at relative trends.

#### 4.3.2 Views per article

We calculate views per article as you’d expect:

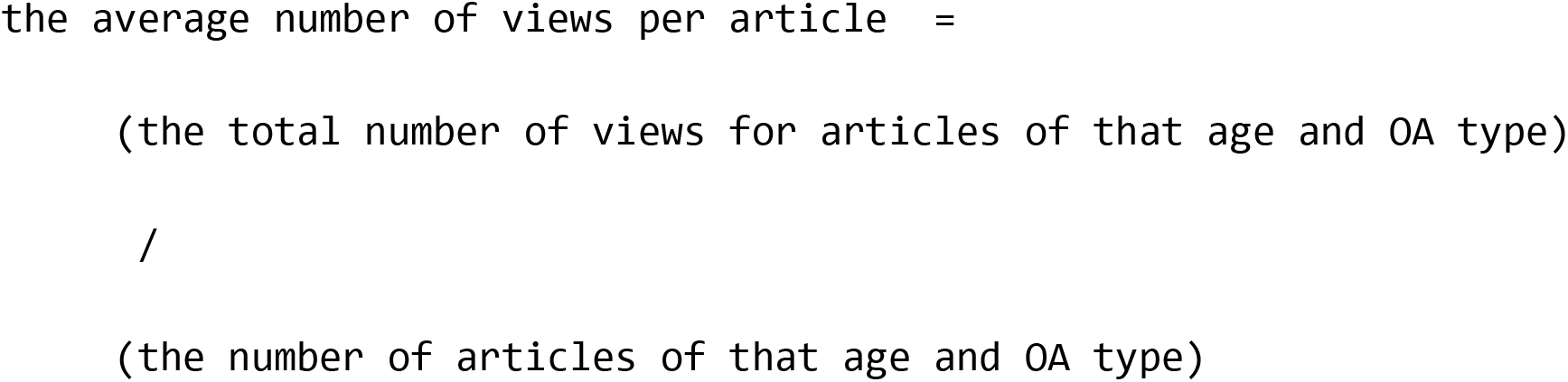

We can state this more concisely and precisely as follows. For each OA type:

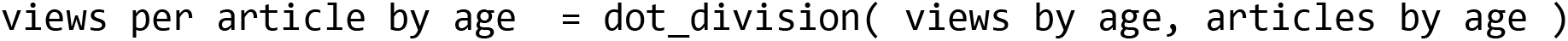

where dot_division is the <u>element-wise Hadamard division</u>) of two signals.

It is important to look at views by article age, because it is well known that readers are more interested in accessing newly-published articles, and indeed this trend can be seen in the Unpaywall usage logs. Figure 13 shows monthly access requests to the Unpaywall extension made between August 2018 and August 2019, distributed by article age. As expected, the distribution is very skewed and readers are most interested in articles published less than a year ago.

**Figure 13:**
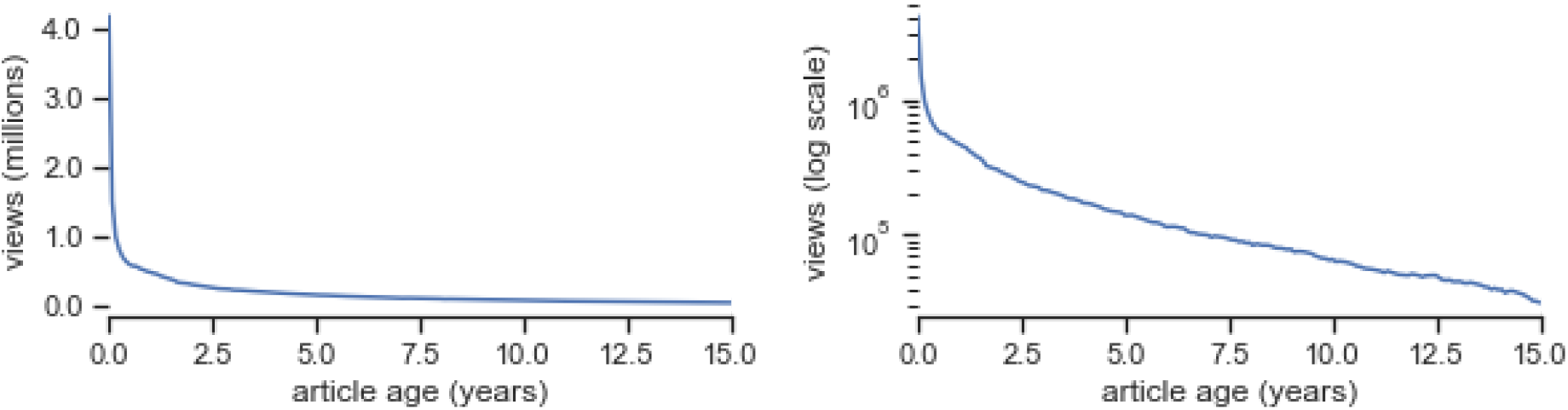
Distribution of views by article age: left panel is linear y-axis and y panel is a log plot.

We now look at this data by OA type. To simplify interactions, for the rest of the analysis we will restrict our view data to that from just one month: July 2019, and to articles less than 15 years old. This accounts for 2,774,403 views, which we then multiply by 12 to approximate a year’s worth of views by the Unpaywall extension (not important because the point of the views analysis is growth rather than absolute numbers, but it helps slightly with interpretation).

Figure 14 shows the distribution of views by article age, by OA type.

**Figure 14:**
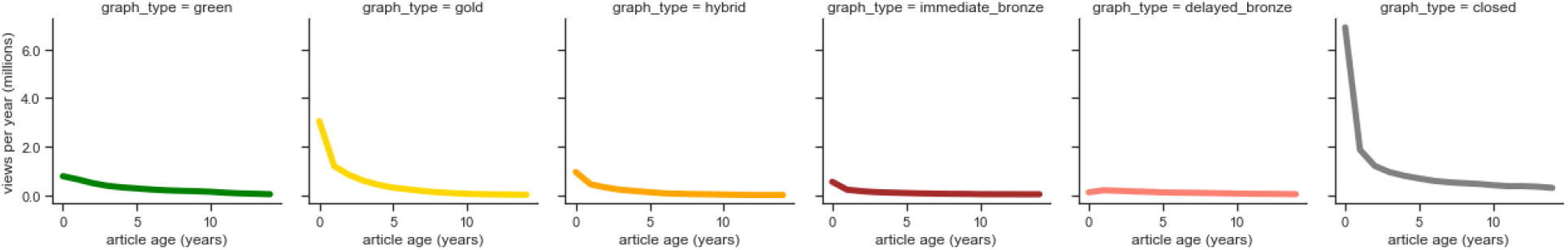
Distribution of views by article age, by OA type.

Closed access articles receive the most views, which isn’t surprising because as we saw in Section 4.1.6, most articles available in 2018 are Closed access. What happens when we divide these curves to get views *per article*, as is our goal?

A detailed walkthrough is given in Supplementary Information Section 11.4. Here we present the results of dividing the above view distribution signals by the article signals we calculated Figure 10 for the 2018 observation year:

We see in Figure 15 that number of views per article is much higher for Green than other kinds of articles, particularly for articles that are available as Green OA within the first year of publication (age 0). This is consistent with previously-documented download advantages of Green OA articles, and could be caused by various factors including self-selection bias, or the common cause of strong funding support for high-interest medical papers by funders like the NIH.

**Figure 15:**
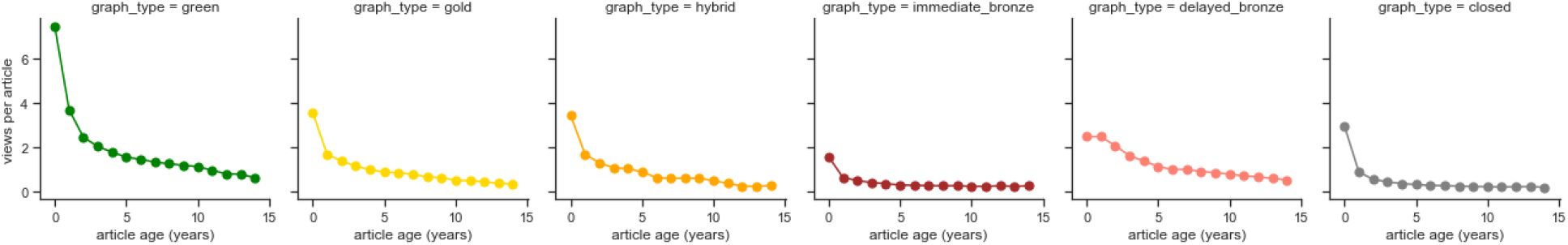
Views by article curve, by OA type.

Relative to Closed access articles, the average number of views per article for Gold, Hybrid, and Delayed Bronze is particularly strong for older articles.

#### 4.3.3 Calculating Views

Finally, we are ready to calculate overall views. As a reminder, here is our general approach:

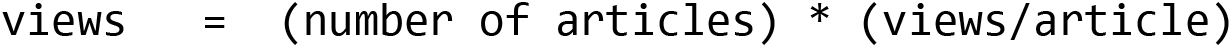

We can state this more precisely as follows. For each OA type:

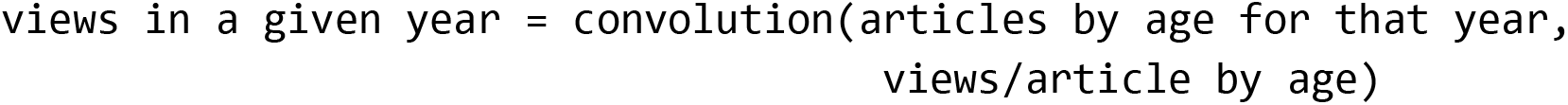

where <u>convolution</u> is the standard mathematical operation of modifying a signal by another signal, by integrating the product of the two curves after one is reversed and shifted.

A detailed walkthrough of this convolution (and more information on what convolution means!) is given in Supplementary Information Section 11.5. Here we present the results of convolution, which can be seen roughly as multiplication of the articles-by-age estimates we made in Section 4.2.2 and Section 4.2.3 with the views/article curves we calculated in Section 4.3.2.

### 4.3.4 Combined Past and Future Views

We can plot these lines stacked on top of each other to see how the OA types change over time, shown in Figure 17.

**Figure 28:**
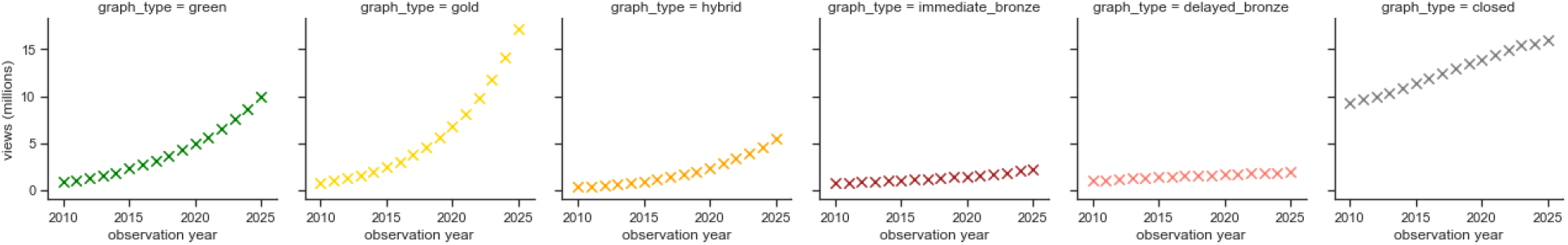
Views. Views by year.

**Figure 17:**
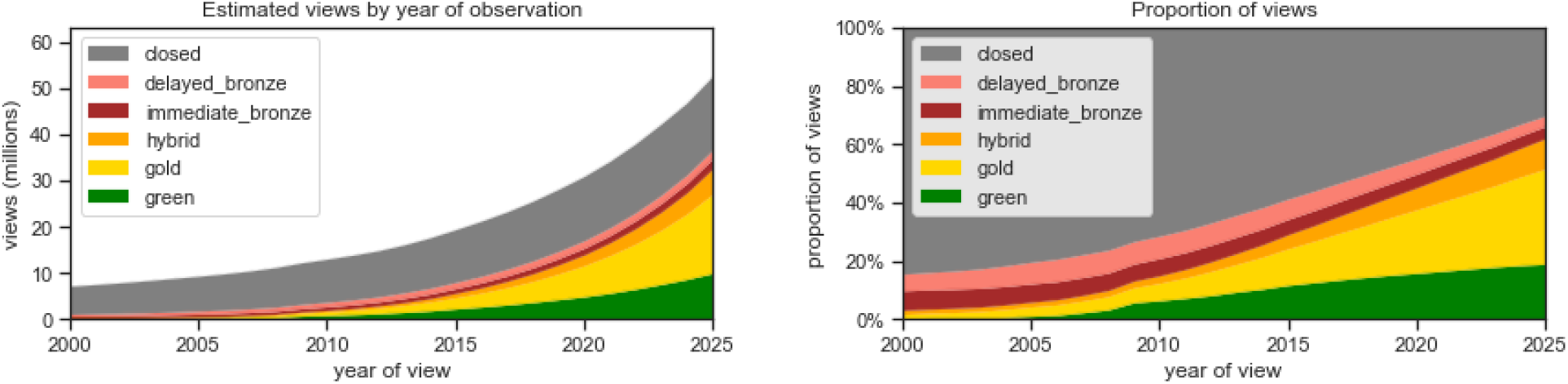
Views, by year of view.

Here is the raw data for the proportions at a few specific observation years:

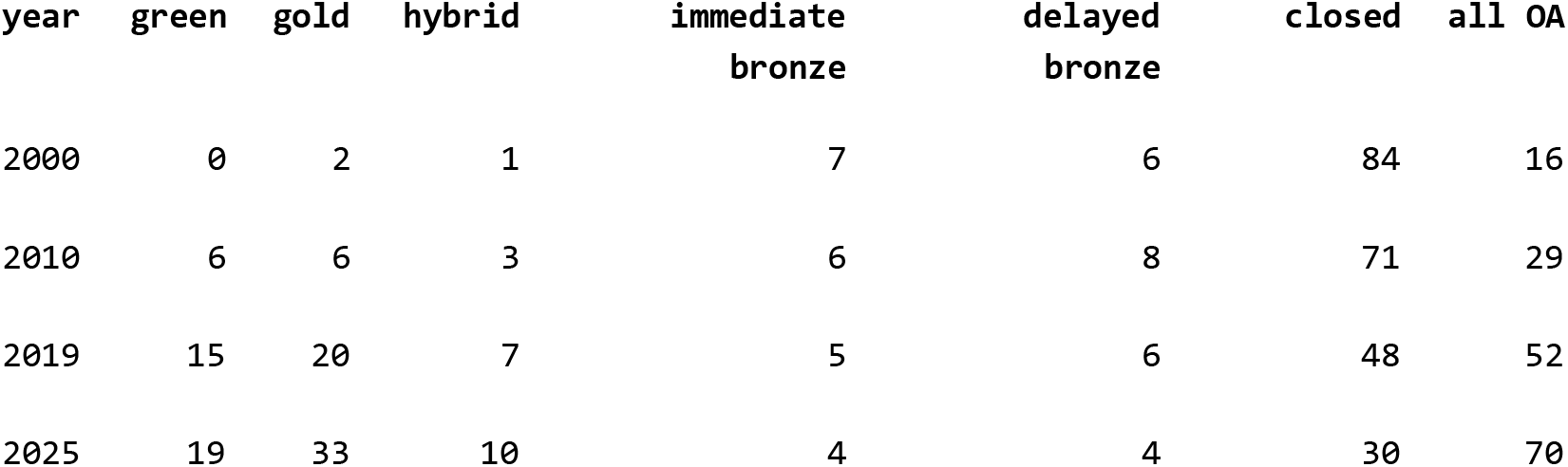

Our estimated number of views per year increases steadily over time, and the proportion of views to OA resources goes from 16% in 2000 to 52% in 2019, and to 70% in 2025. This increase is driven primarily by Green and Gold OA: we estimate 19% of views in 2025 will lead to Green OA articles and 33% of views will lead to Gold articles.

### 4.4 Extending the model: Growth of bioRxiv

An advantage of building a model is that now we can layer on alternate assumptions and see how anticipated disruptions might affect OA in coming years. A comprehensive examination of all the alternative futures is clearly outside the scope of this paper, however an example will be illustrative.

BioRxiv, a preprint server in biology, provides an excellent example. As described in Abdill and Blekhman (2019), deposits into bioRxiv are growing rapidly. If growth continues at the current rate, biorxiv could prove to be a major disruptor: it is growing extremely quickly, and the vast majority of the deposits which are published have zero OA lag (and so are OA at the time of highest demand).

We model the growth of bioRxiv and its impact on OA availability by extrapolating from bioRxiv papers that:

- were deposited at or before the date they were published (to simplify the model), and
- are Closed access other than their Green bioRxiv copy (so that we don’t double-count articles made OA as Gold, Hybrid, or Bronze).

The number of articles that meet these criteria are shown in the table below, by date of publication.

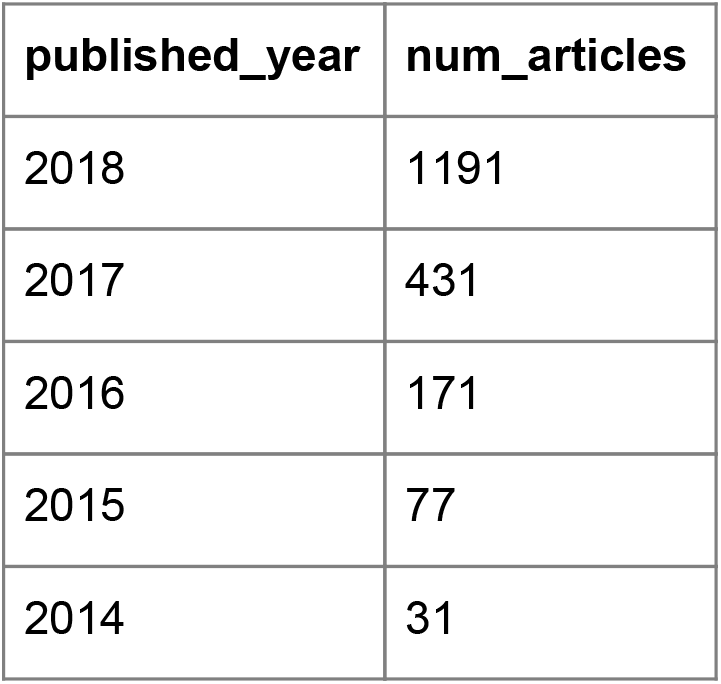

This growth has a very strong logarithmic extrapolation fit, as seen in Figure 18.

**Figure 18:**
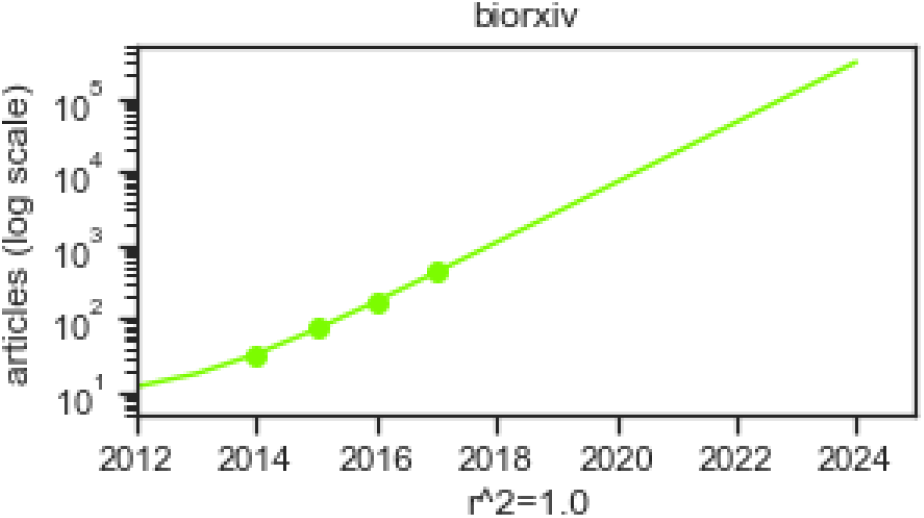
bioRxiv extrapolation.

BioRxiv won’t be able to grow exponentially forever -- there are a limited number of papers in Biology. But if we were to imagine bioRxiv continued its current growth rate for another 5 years, we would estimate its impact on the relative proportion of articles available as OA in Figure 19.

**Figure 19:**
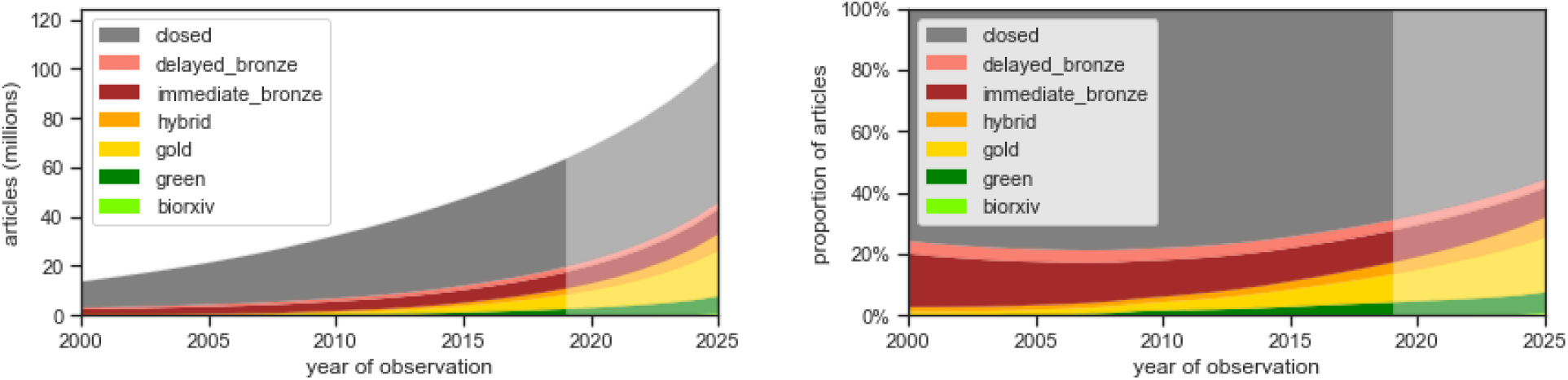
Prediction of articles by OA type by year of observation, including bioRxiv.

This doesn’t look like many articles, but let’s see how it affects viewership. For simplicity we use the generic green OA access trend derived in Figure 24. This results in views as shown in Figure 20 -- a notable impact on the total views of all scholarly papers, and obviously the impact would be even greater within the field of biology.

**Figure 20:**
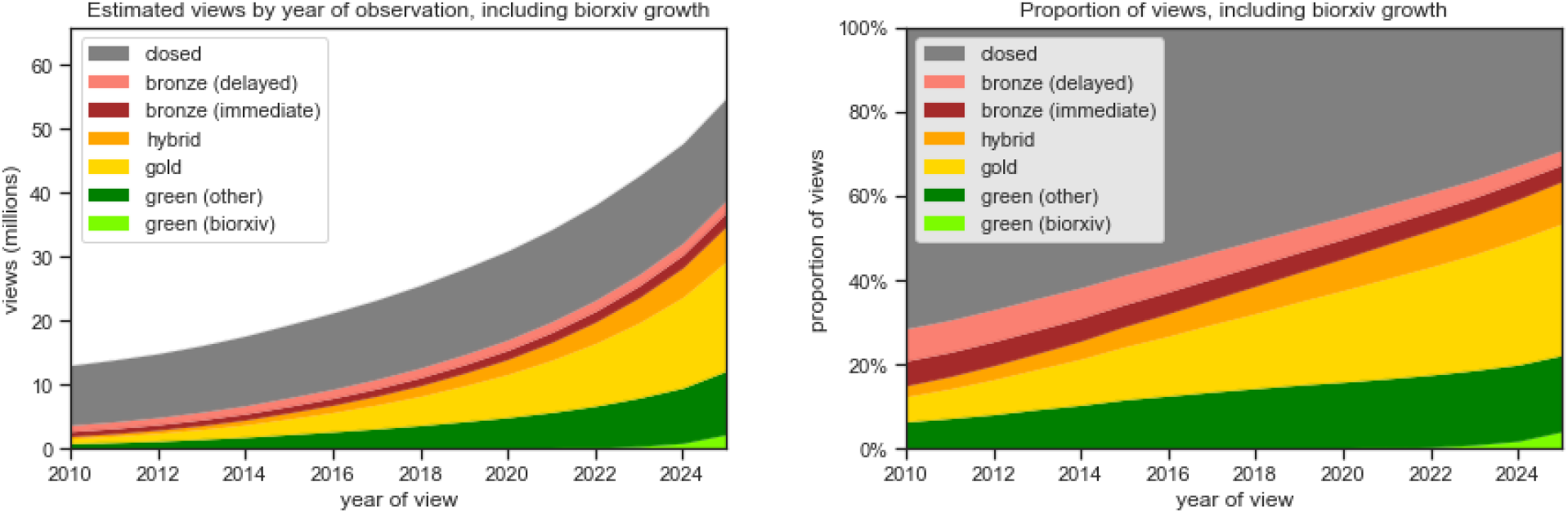
Predicted views, including bioRxiv, by year of view.

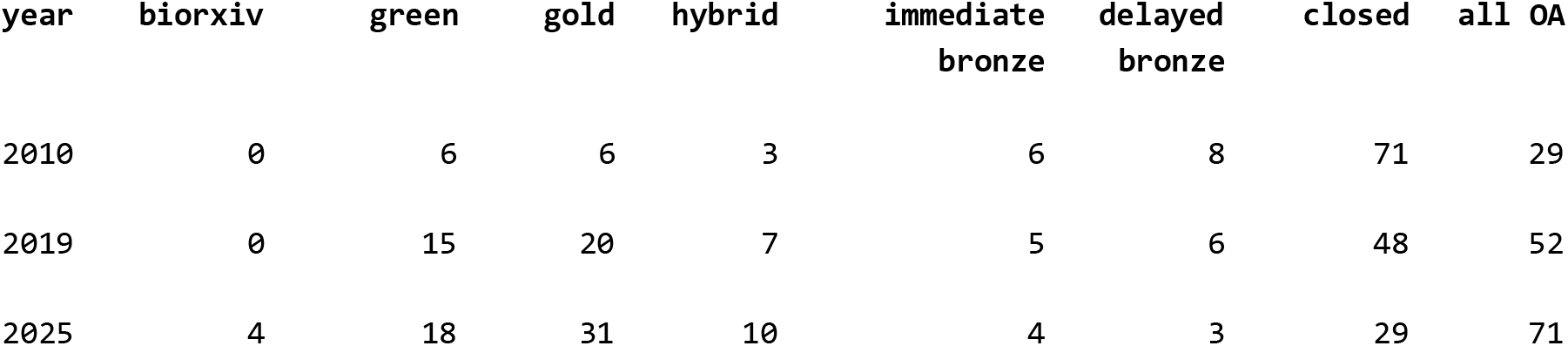

## 5. Discussion

We found that Green, Gold, and Hybrid papers receive more views than their Closed or Bronze counterparts, particularly Green papers made available within a year of publication. We also found that the proportion of Green, Gold, and Hybrid articles is growing quickly.

In 2019:

- 31% of all journal articles are available as OA
- 52% of article views are to OA articles

Given existing trends, we estimate that by 2025:

- 44% of all journal articles will be available as OA
- 70% of article views will be to OA articles

Our model is conservative. Although the extrapolations assume continued incremental adoption of OA, they do not yet model other disruptive changes that will likely increase the growth of OA in coming years: the adoption of Plan S, a change in embargo periods for existing mandates, a dramatic increase in institutional self-archiving, large scale read and publish agreements, etc.

This area is ripe for future research in other ways as well: understanding how OA publication and viewership rates vary by discipline, country, and publisher is key. The assumption that the views/article curve is stable over time should be further investigated and relaxed if found to be inadequate. The model could be refined in many other ways as well, for example to use custom viewership patterns for readers within a specific university, or of a specific journal.

One interesting realization from the modeling we’ve done is that when the proportion of papers that are OA increases, or when the OA lag decreases, the total number of views increase -- the scholarly literature becomes more heavily viewed and thus more valuable to society. This is intuitive, but could be explored quantitatively in future work.

The study has several limitations. Only journal articles with DOIs are included, which under-represents disciplines and geographical areas which rely heavily on conference papers or articles without DOIs. Illegal repositories (SciHub) or articles posted on academic social networks (ResearchGate and Academia.edu) are not considered, which may undercount articles that are relevant for some uses. The users of the Unpaywall browser extension may not be representative of other readers, and using page views as a proxy for article interest is inexact. Nonetheless, we believe this analysis represents a useful approach for modeling the growth and importance of OA in the future.

The genesis for this study was a steady stream of inquiries from university librarians, asking for OA rates for specific journals to help inform their subscription decisions and negotiations. We realized it would be even more helpful if we could provide OA rates (a) for the future, (b) by date when the OA resource is available, and (c) weighted by the importance of the article to their faculty. The model presented here addresses these issues and will form the basis of information available to librarians and other decision-makers in the future.

## 7. Data and code availability

### 7.1 Empirical Gold OA list

The empirical Gold OA journal list is available in the Zenodo dataset at http://doi.org/10.5281/zenodo.3474007, in the file “gold_oa_empirical_list.csv”.

### 7.2 Empirical bronze delayed OA list

The empirical Bronze Delayed OA journal list is available in the Zenodo dataset at http://doi.org/10.5281/zenodo.3474007, in the file “delayed_bronze_empirical_list.csv”.

The list of combined delayed OA policies we extracted from various sources is available in the Zenodo dataset at http://doi.org/10.5281/zenodo.3474007, in the file “delayed_bronze_extracted_policies.csv”.

### 7.3 Study data

All study data is available in Zenodo, at Piwowar H, Priem J, & Orr R. (2019). Data From: The Future of OA: A large-scale analysis projecting Open Access publication and readership [Data set]. Zenodo. http://doi.org/10.5281/zenodo.3474007

### 7.4 Analysis notebook

The Jupyter analysis notebook is available from GitHub at https://github.com/Impactstory/future-oa.

Also, for Jupyter nerds and to help us remember: export using jupyter nbconvert manuscript.ipynb --to html --TemplateExporter.exclude_input=True then push to github, then can be viewed at https://htmlpreview.github.io/?https://github.com/Impactstory/future-oa/blob/master/manuscript.html

## 10. Acknowledgements

The authors would like to thank Bianca Kramer for extensive and valuable comments on a draft of this article. The author order of JP and HP was determined by coin flip, as is their custom.

## 11. Supplementary Information

### 11.1 Detailed look at OA Lag of Green OA

This data supplements the discussion of Green OA lag in Section 4.1.2.

In Figure 21 we plot the number of Green OA papers made available each year vs their date of publication. The first plot is a histogram of number of papers made available each year (one row for each year). The second plot is the same, but superimposes the articles made available in previous years. This stacked area represents the total cumulative number of Green OA papers that are available in that year -- if you were in that year and wondering what was available as Green OA that’s what you’d find.

**Figure 21:**
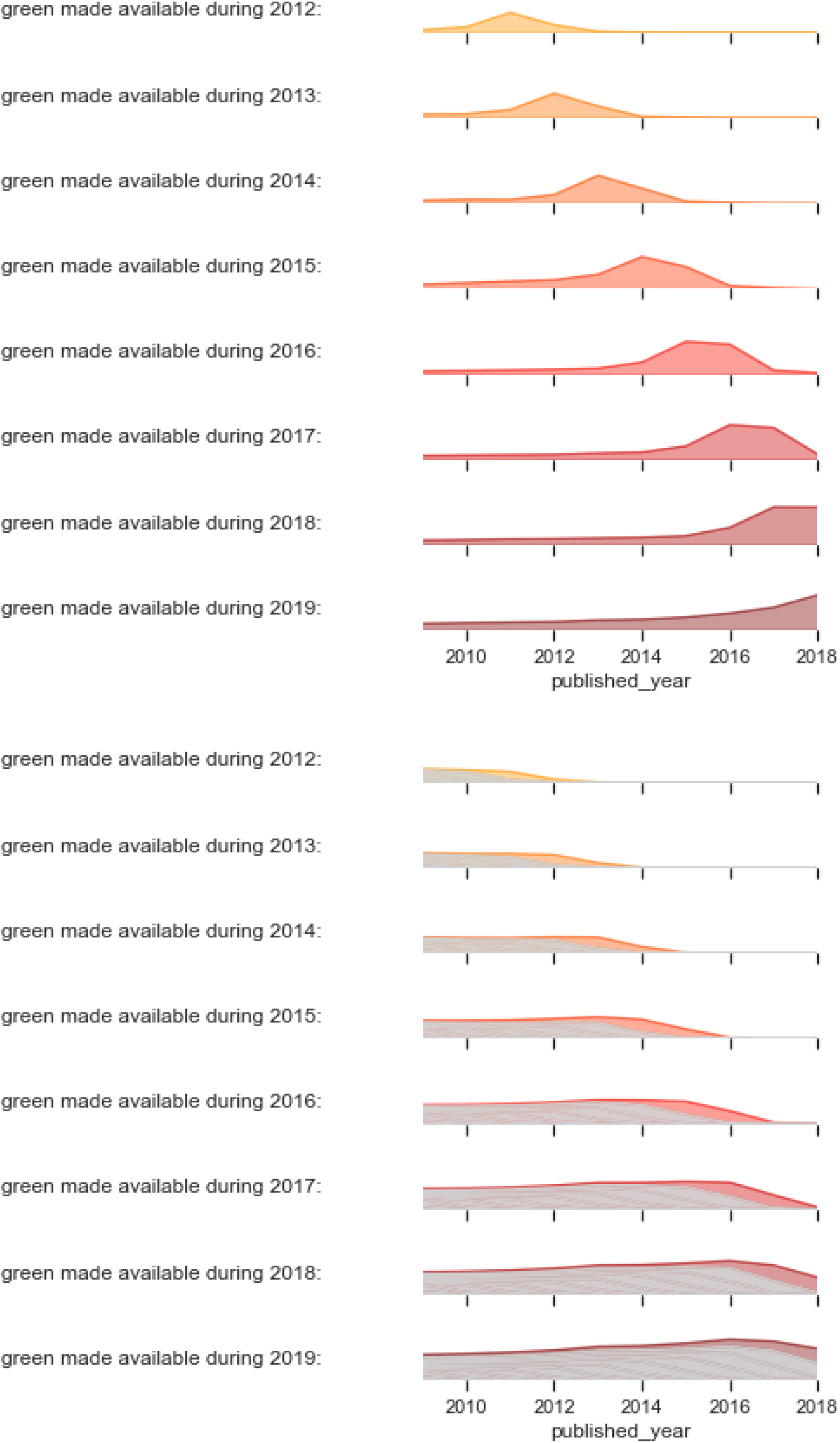
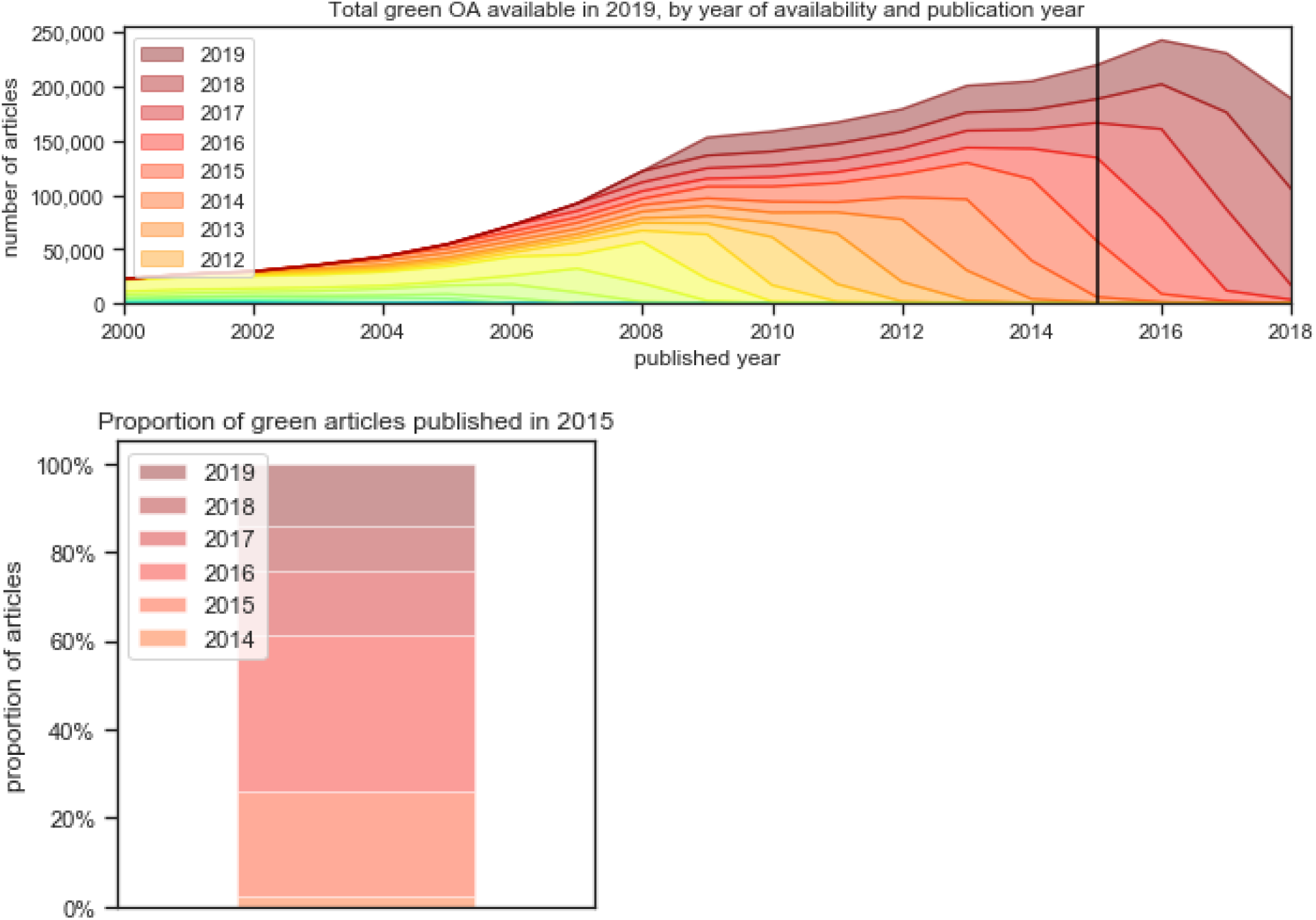
Detailed Green OA.

The third plot is a larger version of the availability as of 2018, showing the accumulation of availability. It allows us to appreciate that less than half of papers papers published in, say, 2015, were made available the same year -- most of the papers have been made available in subsequent years. The fourth plot is a slice in isolation, for clarity: the Green OA for articles with a Publication Date of 2015.

### 11.2 Detailed look at OA Lag of Delayed Bronze OA

This data supplements the discussion of Delayed Bronze OA lag in Section 4.1.3.

In Figure 22 we plot the number of Delayed Bronze OA papers made available each year vs their date of publication. For more explanation see the text describing Figure 21 above.

**Figure 22:**
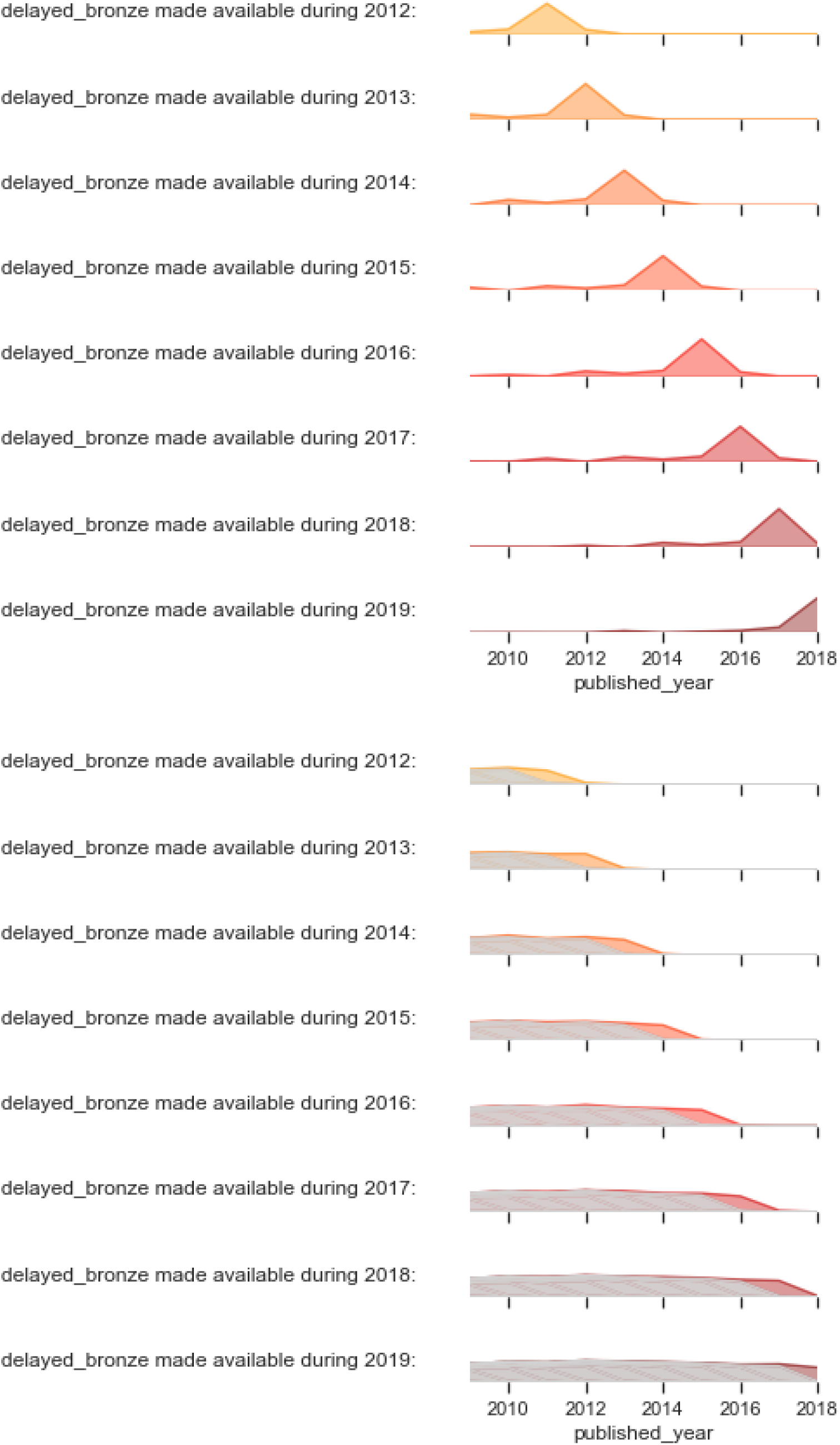
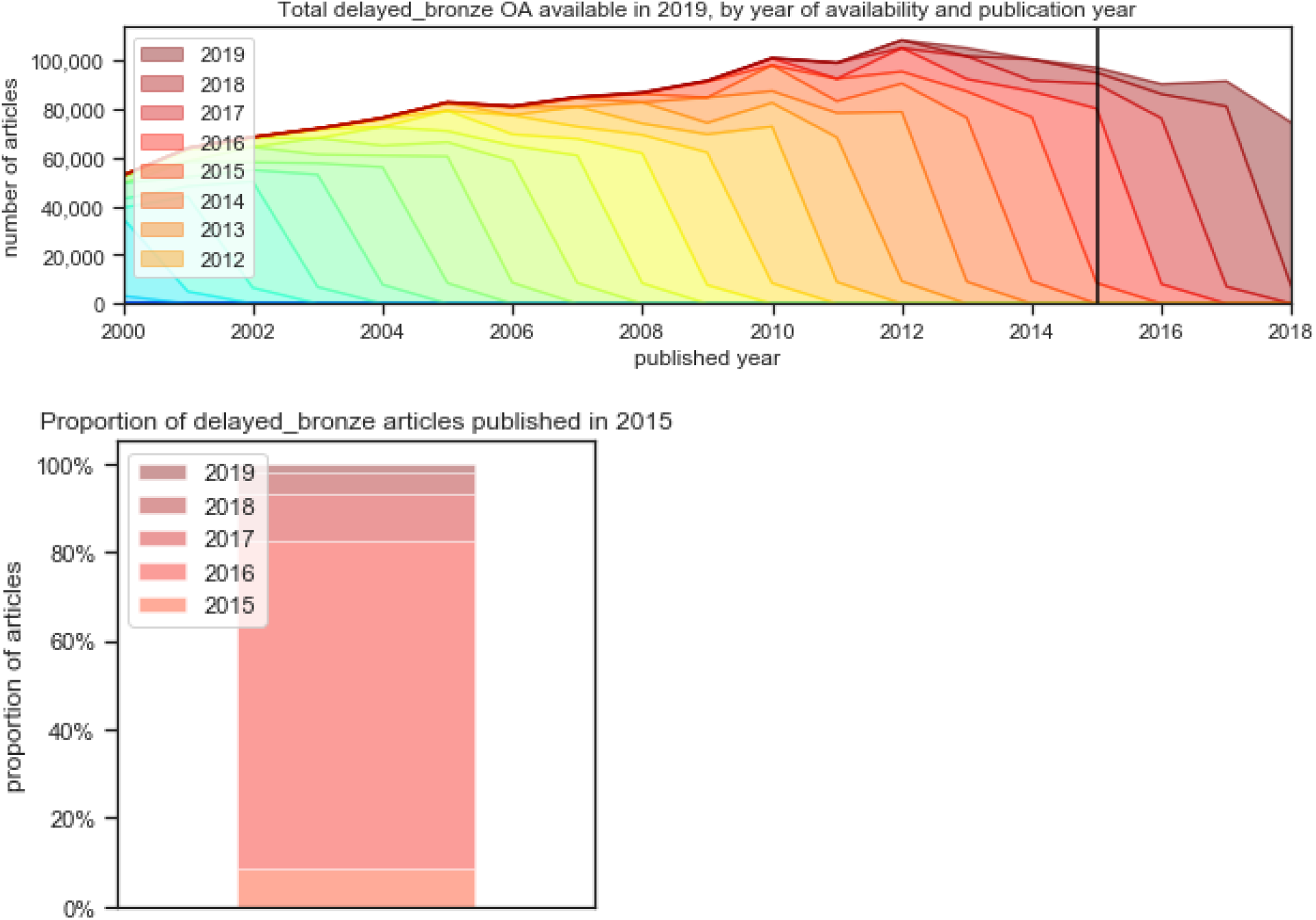
Detailed Delayed Bronze OA.

### 11.3 Walk-through of dot division

This is a walk-through of dot division, as discussed in Section 4.3.2. For each OA type:

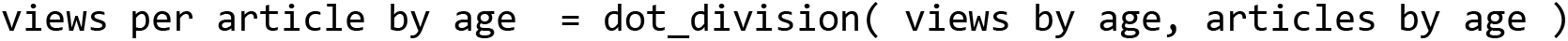

where dot_division is the <u>element-wise Hadamard division</u>) of two signals.

For **views by age** we use Figure 14, reproduced here:

**Figure 14:**
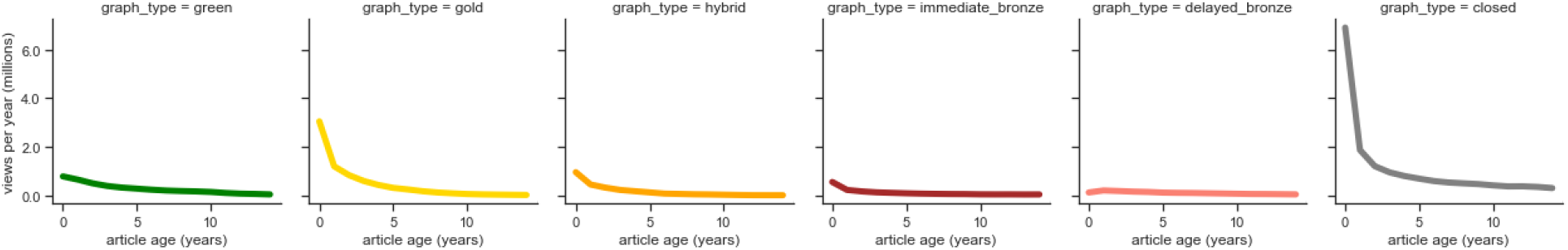
Views by OA type and article age.

For **articles by age** we use the curves we calculated in Section 4.2.7 above, specifically Figure 10 for the 2018 observation year:

**Figure 23:**
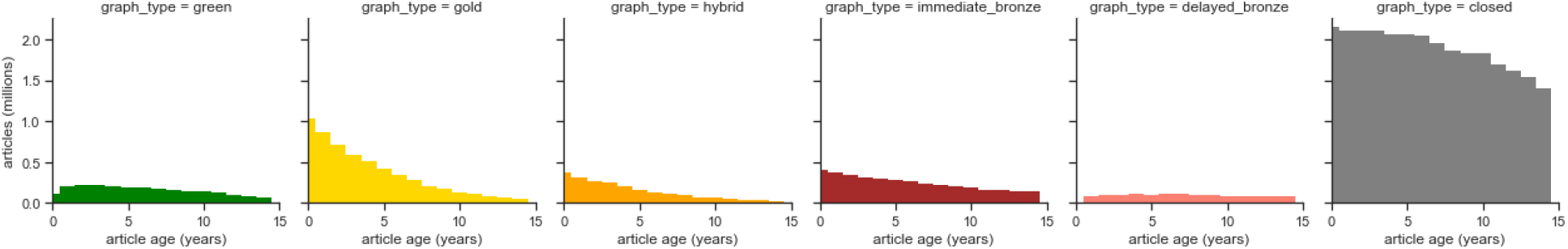
Articles by OA type and article age.

Now for each of the OA types individually, we divide these signals by each other, element-wise. This means that, for each OA type, we divide the number of times someone viewed an article of that was 0 years old (in other words, published less than a year ago) by the number of total articles that were 0 years old -- articles that were published less than a year ago. Then we take the next age bucket, 1 years old, and divide the number of views of 1 year old articles by the number of articles available as that OA type that were 1 year old. We do this for all age bins (15 years are shown in the graphs).

The result of these divisions are the signals below: the number of views per article, for a given age and OA type.

**Figure 24:**
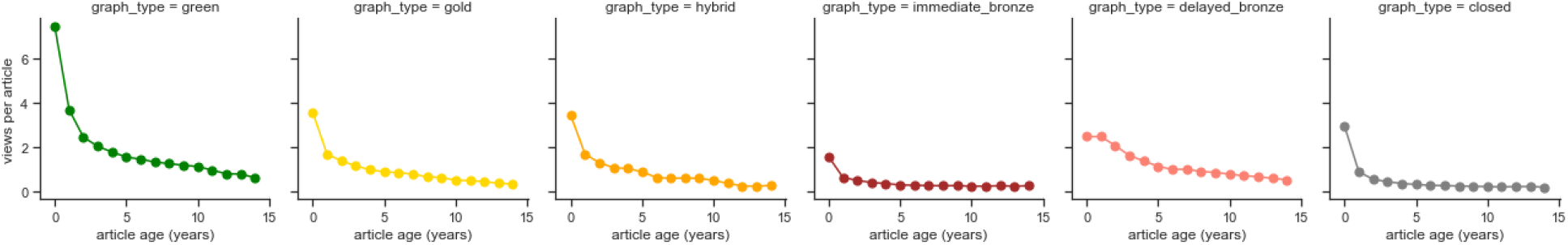
Views by article curve, by OA type.

### 11.4 Walk-through of convolution

This is a walk-through of convolution, as discussed in Section 4.3.3. For each OA type:

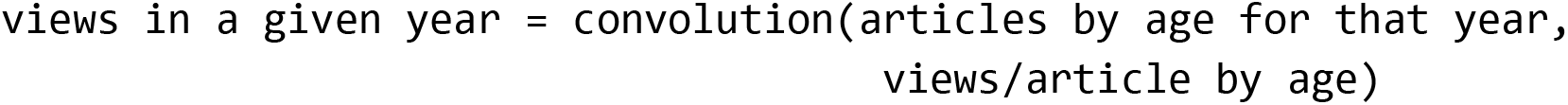

As an example, an estimate of total number of views in 2022 comes from summing together the views in 2022 across all article ages. In other words, the green X in the graph below at year 2022 is the area under the green curve in the row above -- the sum of all views to green OA of age 0, age 1, age 2, age 3, etc. We then did this for all years.

To show how we’ll estimate views, we’ll use 2022 as an example. We use the 2022 row from above, and the graph it by age of article (rather than year of publication). This flips the direction of the x axis. In this graph to make the next steps more clear we also use a shared y axis across all OA types.

**Figure 25:**
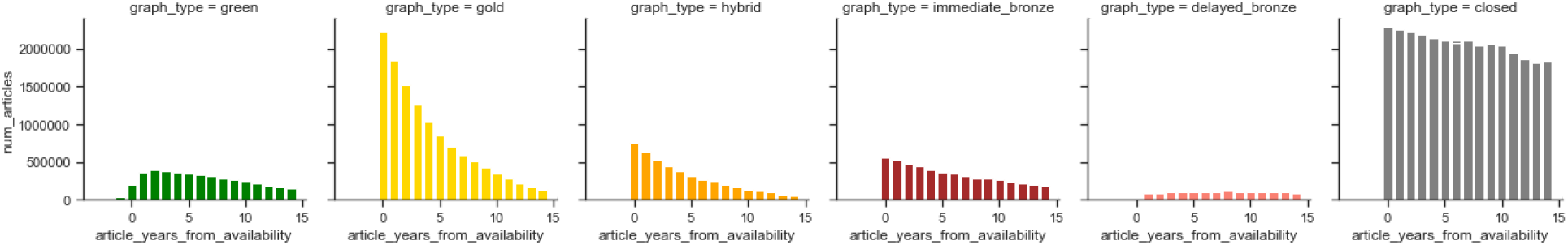
Predicted number of articles by OA type by article age, at the year of observation 2022.

Next, we’ll use the signal we calculated in the section “How often does someone want to access a paper, given its age and OA type”, which shows the number of views per article someone made in 2019. An assumption in our model is that this views-per-article probability stays the same across time, so we assume that it applies to 2022 as well.

**Figure 26:**
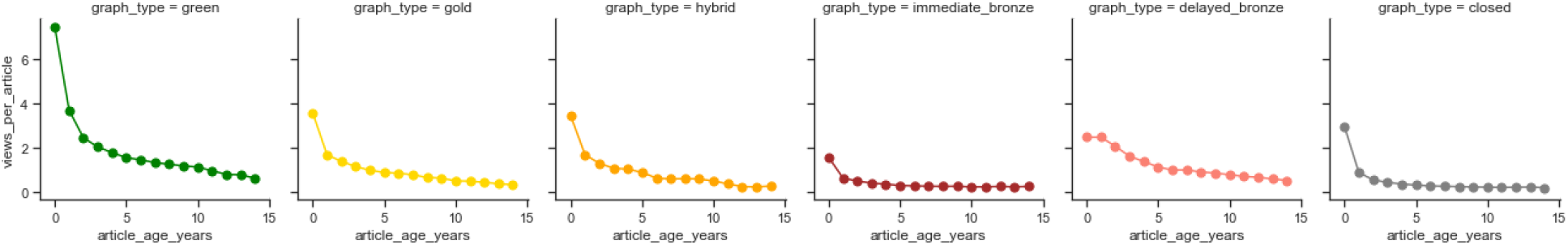
Views per article by OA type by article age.

Now we multiply these two signals together. We multiply them in a similar way that we divided signals in an earlier step -- we take each OA type in turn, and then take each age bin in turn. So the green OA point at 0 years in the graph below comes by multiplying the number of estimated articles in 2022 that are available as green OA and 0 years old by the number of “views-per-article” we calculated for green OA for articles that are 0 years old. We then do that calculation for every age bin, for every OA type, and get the graph below:

**Figure 27:**
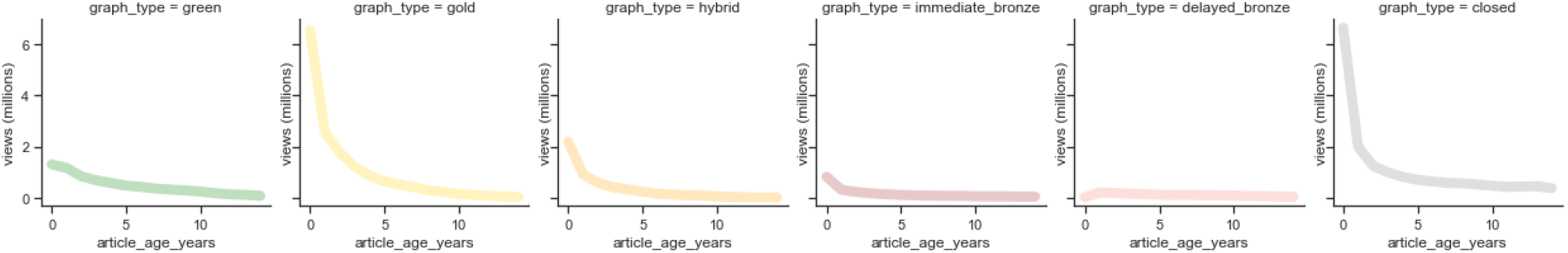
Views by OA type by article age for the observation year 2022.

**Figure 28:**
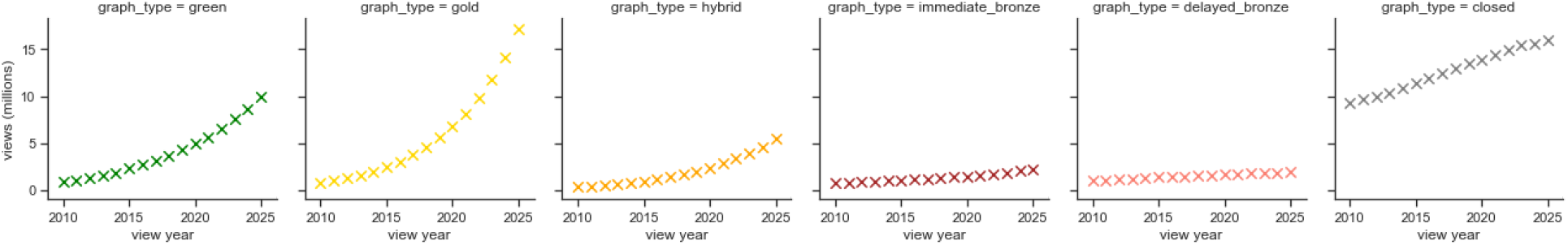
Views. Views by year.

This gives us views for each year, 2010 to 2025, by OA type. The following graph is the same as the previous one, but without shared y axis so we can better see the relative trends.

**Figure 29:**
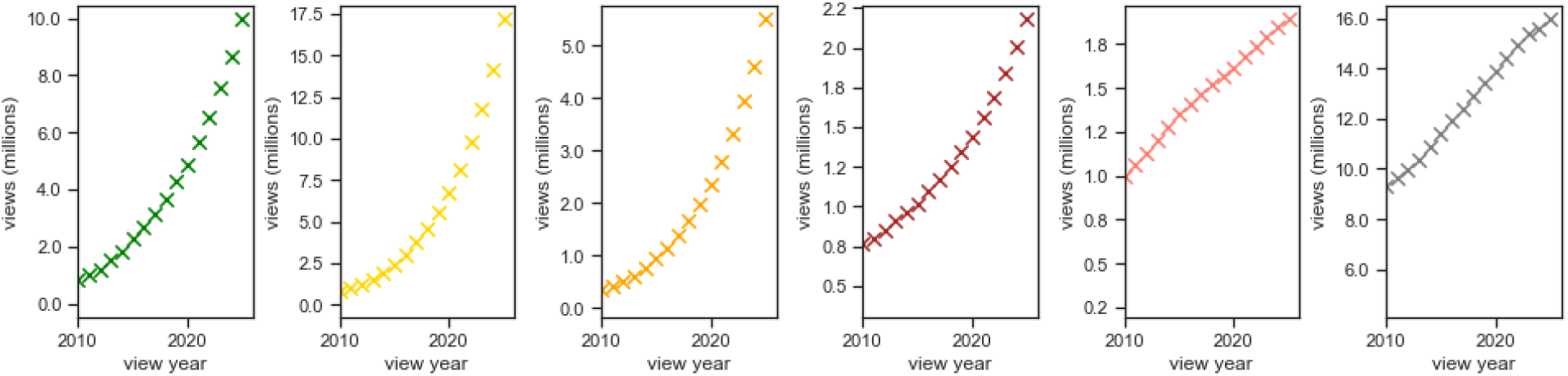
Views by year, full scale.

### 11.5 Number of articles by OA type, by date of publication

You may be wondering what the data would look like if we ignored the idea of year of observation and simply plotted articles by year of publication, categorized by OA type as we measure it today. We present this data in Figure 30.

**Figure 30:**
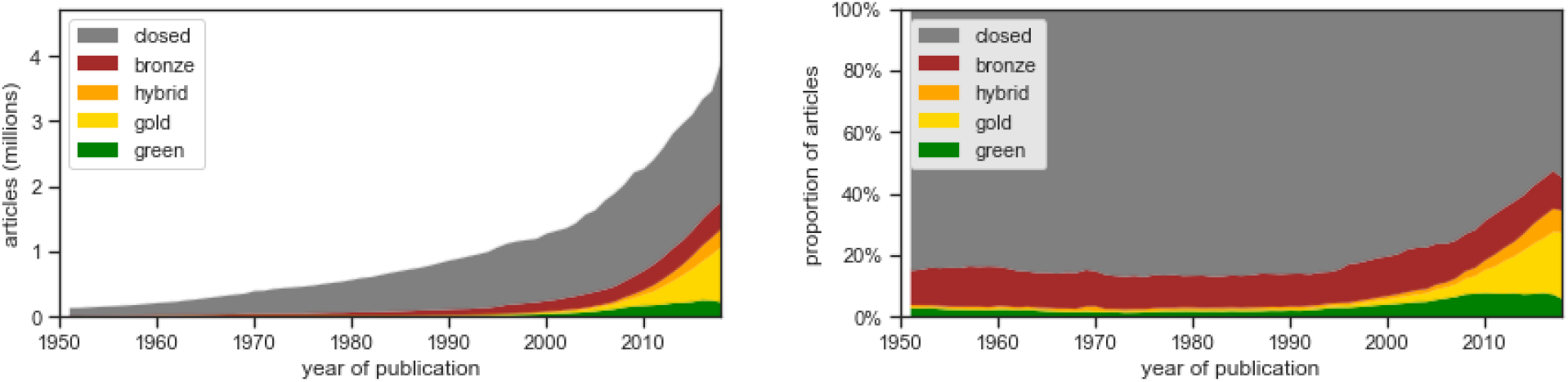
Articles by OA type, by year of publication. OA type as of October 2019.

The early years of Figure 30 is similar to Figure 2 in Piwowar et al. (2018), with one notable difference: some of what was considered Bronze OA (and to a lesser extent hybrid OA) in Piwowar et al. (2018) is classified as Gold OA in the current analysis. This is due to an improvement in Unpaywall’s algorithms. Originally, Unpaywall used the Directory of Open Access Journals (DOAJ) as the sole arbiter of whether a journal was “fully-OA.” Unpaywall still uses DOAJ in this way, but it now also adds an empirical check for OA journals (if 100% of a journal’s articles are OA, it is listed as an OA journal). This results in a more comprehensive and accurate list of fully-OA journals, which in turn moves some articles into Gold from Hybrid and Bronze. We’ve made this comprehensive Gold OA journal list available: see Section 7.1.

We can see a visible decrease in the proportion of OA (particularly Green) in the most recent publication years. **This change in OA proportions is because many articles published in 2018 are still under embargo at the time of this analysis (October 2019), so they are considered “Closed” in this graph even though they may ultimately become Green OA or Bronze OA.** More on this in Section 4.1.

A cumulative view of Figure 30 is shown in Figure 31.

**Figure 31:**
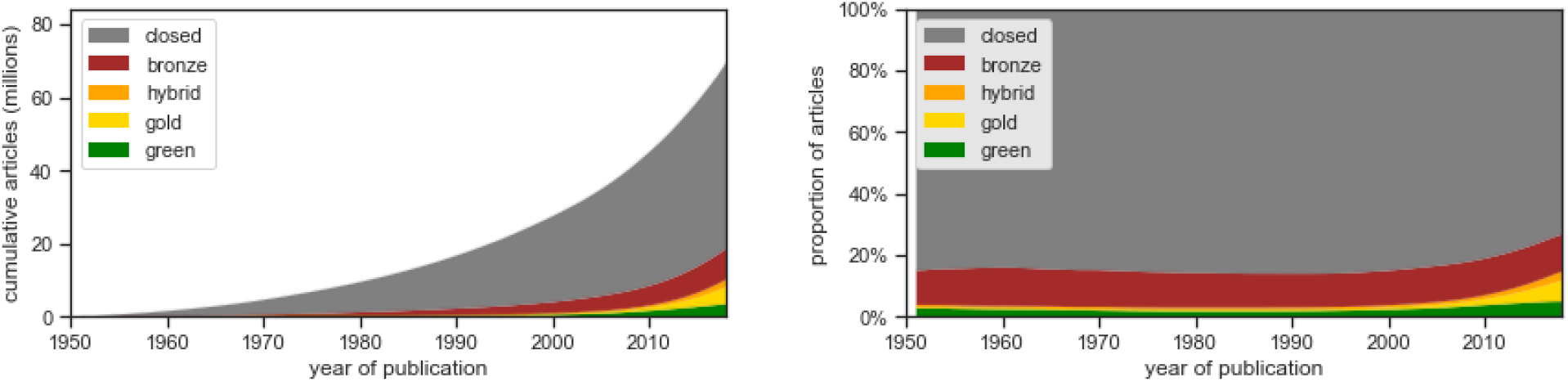
Cumulative articles (total articles extant in the world) by OA type, by year of publication. OA type as of October 2019.

